# Evaluation effects can rival reported concordance gains in cross-cancer survival prediction

**DOI:** 10.64898/2026.11.05.764739

**Authors:** Ricky Nguyen, Fatemeh Vafaee

**Affiliations:** School of Biotechnology and Biomolecular Sciences, University of New South Wales (UNSW Sydney), Australia; UNSW AI Institute, University of New South Wales (UNSW Sydney), Australia

**Author notes:** Correspondence to: Professor Fatemeh Vafaee, School of Biotechnology and Biomolecular Sciences, UNSW SYDNEY NSW 2052 AUSTRALIA, E.

**Keywords:** multi-omics, survival prediction, cross-cancer transfer, model evaluation, model selection, external validation, reproducibility

## Abstract

Studies of cross-cancer learning for survival prediction from tumour omics have reported concordance gains of 0.020 and 0.045, but how such gains compare with evaluation effects remains unclear. We examined these effects across 13 TCGA cancers (5,761 patients, 1,526 disease-specific deaths), using five repeats of five-fold cross-validation and paired bootstrap intervals. When model selection did not use test data, our borrowing model gained 0.009 (95% interval 0.001 to 0.016) over its target-fitted reference when cancers received equal weight (post hoc). Random-effects pooling gave a gain of 0.002 (−0.002 to 0.006), with most weight assigned to cancers where the model made little use of the pretrained score. Choosing the fusion weight on test splits raised the apparent equal-weight gain to 0.017. We reimplemented a published method, VAECox, which showed no pretraining gain over random initialisation at the last or validation-selected training step. Choosing each network’s best test step, as the published code does, raised concordance by 0.03 to 0.05 across the 13-cancer and ten-cancer overall-survival settings, but barely changed the difference between networks. In a constructed post hoc experiment, choosing the best test step only for the pretrained network changed the equal-weight mean difference from −0.004 to +0.045 (+0.035 to +0.054). Across seven multi-omics comparisons, estimates from single five-fold runs differed from their cancer’s five-run mean by at least 0.02 in 11% to 42% of runs. Concordance was about 0.02 lower at held-out tissue source sites. Interpreting a reported gain therefore requires each model’s selection rule, variation across repeated partitions, the weighting of cancers and the evaluation population.

**Key Points:**

- In a constructed post hoc experiment, applying test-step selection to one of two VAECox networks only shifted their mean difference by 0.027 to 0.049, comparable to reported gains of 0.020 and 0.045, and single five-fold runs deviated from their cancer’s five-run mean by at least 0.02 in 11% to 42% of runs across seven multi-omics comparisons.
- Evaluating both VAECox networks at their best test steps raised absolute concordance by 0.03 to 0.05 but changed their mean difference by about 0.002 or less. Holding out tissue source sites lowered concordance by about 0.02; whether the decrease differed between the two models evaluated was inconclusive.
- With test-independent selection, the borrowing gain was +0.002 under the random-effects summary, which was dominated by cancers in which little borrowing occurred, and +0.009 with equal weight per cancer. A reimplementation of VAECox showed no average pretraining gain with the published-style procedure, with the target cancer excluded from pretraining, or with the fine-tuning step chosen on validation data.
- A reported concordance gain should be accompanied by how each model’s hyperparameters and training length were selected, whether the models compared were treated alike, the variation across repeated partitions, how cancers were weighted and which patients or sites were held out.

## 1 Introduction

Bulk RNA, microRNA and DNA methylation profiles, treated here as three omics blocks, are available for thousands of tumours [1, 2], and many models have been proposed to use these data for survival prediction [3–5]. A recurring idea is that a model for a given cancer should borrow information from other cancers through pan-cancer pretraining or joint training, because individual cohorts typically contain only a few hundred patients and far fewer events [6–8]. Many such models are deep multimodal architectures that learn a shared representation of the omics blocks [9]. Two studies of cross-cancer learning that trained on cohort-scale target data, not a few-shot subset, reported improvements in concordance of 0.020 [10] and 0.045 [11] over models trained on the target cancer; the first pretrains on RNA and the second trains jointly on histology, RNA and clinical text, and they also differ in reference models and evaluation design (Supplementary Note 1).

Three large benchmarks have examined aspects of these ideas. Herrmann et al. [3] compared eleven statistical methods across 18 TCGA cancers and found that BlockForest was the only method to outperform a clinical-only Cox model, although the difference was not statistically significant. Wissel et al. [4] expanded the comparison to include neural models and found that BlockForest, PriorityLasso and mean late fusion performed best. SurvBoard [5] extended the evaluation to 28 datasets and reported that joint pan-cancer training improved median performance for most methods. All three benchmarks evaluated overall survival. Nießl et al. [12] showed that choices of datasets, performance measure and aggregation change which method appears best. Here we ask how large such effects are on a paired gain from cross-cancer learning, as distinct from absolute concordance.

As an illustrative audit, we examined twelve selected peer-reviewed studies of cross-cancer survival learning [7–11, 13–19] (Supplementary Note 1). Their evaluation designs differ in target-cancer inclusion, target training size, resampling and uncertainty reporting. These designs address different questions, including transfer to a full-size target cohort, adaptation from a few target patients and joint training. The released code of two of them, VAECox [10] and TMO-Net [9], reports the concordance at the best test-fold step. An earlier study by Gross et al. [20] pretrained an autoencoder on TCGA RNA-seq with the downstream cohorts excluded, tuned both arms with equal budgets and found no significant improvement in survival prediction.

Gains of this size are hard to estimate. Across the 13 cancers studied here, test folds contain between 14 and 54 disease-specific deaths (Supplementary Table S1), and concordance estimated from so few events varies by several hundredths between splits. We tested how choices documented in the audited studies, including selection of the training epoch on test data, reporting a single split or a single five-fold run and inclusion of the target cancer’s patients in pretraining, affected absolute concordance and differences between models, and reran a published borrowing method with and without them.

We did this in 13 TCGA cancers under a paired, repeated design with the target cancer excluded from pretraining (Figure 1). We first estimate the gain from borrowing when selection is independent of the test data, for our own configuration and for a reimplementation of VAECox [10]. We then measure how much evaluation choices change such estimates, separating effects on a difference between models from effects on absolute concordance. To check that the reference model for the borrowing analysis, a product-of-experts variational autoencoder (PoE-VAE), is competitive, we compare it with late fusion of per-block models and with the methods ranked highest by the benchmarks; we also evaluate the models in an external cohort and against a model based on up to four clinical variables. We compare the size of these effects with the published gains of 0.020 and 0.045 to identify what must be reported for a gain of a few hundredths to be interpretable.

**Figure 1.**
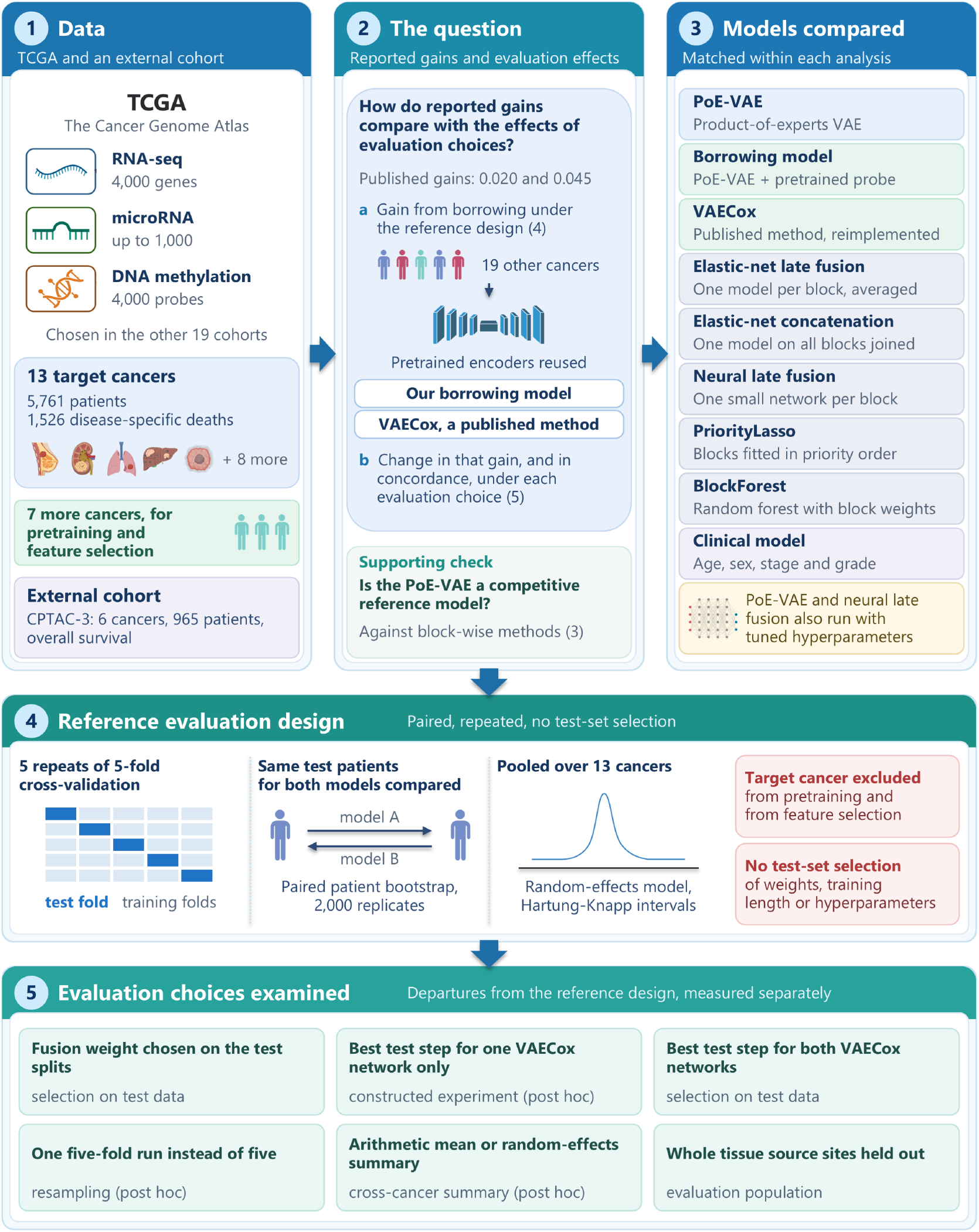
Study overview. (1) Data: RNA, microRNA and DNA methylation from 13 TCGA target cancers, seven further TCGA cohorts used only for pretraining and feature selection, and six CPTAC-3 cohorts for external evaluation. (2) The question: how the published gains of 0.020 and 0.045 [10, 11] compare with the effects of evaluation choices, examined for two borrowing implementations, with a supporting check that the reference model is competitive. (3) The models. The principal molecular comparisons share features, patients and partitions; the VAECox and clinical analyses are separate. (4) The reference evaluation design of the principal analyses. (5) Departures from this design whose effects were measured separately (Section 3.4); three were post hoc. PoE-VAE, product-of-experts variational autoencoder.

## 2 Methods

### 2.1 Data

RNA-seq (STAR TPM), microRNA and 450k methylation matrices and the pan-cancer clinical table of Liu et al. [6] were obtained from the GDC Xena hub [2, 21]. Primary tumour samples were used. A patient was eligible for a cancer’s analysis if they had at least one omics block and a valid disease-specific survival time and status. The 13 targets are the TCGA cancers with at least 75 disease-specific deaths among eligible patients; COAD, UCEC, CESC, ESCA, MESO, KIRP and KICH were used only for pretraining and feature selection. A block was used for a cancer if at least 50 patients had it.

RNA and microRNA values were used on the scale supplied by the hub, log2(TPM + 1) and log2(RPM + 1), without further transformation. Methylation beta values were clipped to [10^−6^, 1 - 10^−6^] and converted to M-values as log2(beta / (1 - beta)) [22]. Imputation and standardisation were fitted on the training patients of each partition (Section 2.2).

### 2.2 Features and splits

Features were selected in the 19 non-target cohorts only. After quality filters applied within each cohort (Supplementary Note 2), features were ranked by the mean of their cohort-specific median absolute deviations, and the top 4,000 RNA, up to 1,000 microRNA and 4,000 methylation features were retained. Missing values within observed blocks were imputed using the corresponding training medians, after which features were centred and scaled using the training patients’ means and standard deviations. This shared feature set was used for the principal internal comparisons of the molecular models; the alternative of selecting features in the target cancer is defined, with its results, in Supplementary Note 2.

For each cancer, we performed five repeats of five-fold cross-validation using different random seeds, with folds stratified by the joint event status for overall, disease-specific and progression-free survival. Each repeat (one five-fold run) comprised five train–test partitions, giving 25 partitions per cancer, and each eligible patient was tested once per repeat.

### 2.3 Models

The reference model, a product-of-experts variational autoencoder (PoE-VAE), was fitted on the target cancer. The borrowing model combined its risk score with a Cox score fitted on the target cancer using representations pretrained in other cancers and then frozen. Both models used the same features, selected in the non-target cohorts. Their comparison therefore measures the added value of the pretrained representation and its fusion with the reference score. We chose the PoE-VAE because its encoders provide the shared representation used for borrowing and can combine any subset of a patient’s observed blocks without imputing an absent block. The architecture was fixed before the comparisons reported here.

PoE-VAE. This model combines the available omics blocks into a shared representation for survival prediction. It uses a product-of-experts variational autoencoder [23, 24] with a proportional-hazards survival head [25]. Each omics block has an encoder with layer normalisation [26]. Each encoder produces a Gaussian posterior over the shared latent variable; the posteriors for the observed blocks are combined with a standard normal prior by a product of experts [24]. A linear projection of the latent variable gives the log-risk, with a piecewise-exponential baseline hazard [27].

The training loss includes the survival likelihood, integrated over the Gaussian log-risk by Gauss–Hermite quadrature [28], the reconstruction error for observed blocks and the Kullback–Leibler divergence [29] from the prior. At each step, the encoders receive a random non-empty subset of each patient’s observed blocks, but the model reconstructs all observed blocks. This follows subset training in multimodal variational autoencoders [24]. Training used 100 full-batch steps of AdamW [30]; concordance was computed from the mean log-risk.

The primary analyses used one fixed configuration, with hyperparameters set in earlier work on KIRC and HNSC. We later added a version with hyperparameters selected by inner cross-validation as a check (described below; results in Section 3.6). Unless labelled “tuned”, PoE-VAE refers to the fixed configuration. Supplementary Note 2 gives the layer sizes, loss and optimiser settings.

#### Pretrained probe and borrowing model

To learn a representation from other cancers, we pretrained the same architecture for 300 full-batch steps on the 19 non-target cohorts. Pretraining used the same objective, a shared risk projection and a separate baseline hazard for each cohort. We then froze the encoders and fitted a ridge Cox model [31] to the latent means of the target cancer’s training patients. This model is the pretrained probe. Its penalty was selected from 10^−2^ to 10^3^ by inner five-fold concordance, with ties favouring the larger penalty. The borrowing model combines the standardised PoE-VAE risk score, *r*_PoE_, and the standardised probe risk score, *r*_probe_, as

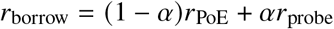

where the scalar fusion weight *α* ∈ {0, 0.1, …, 1} is selected by inner five-fold cross-validation [32], with ties favouring the smaller *α*.

Selection proceeded in two stages within the outer-training set. First, the probe penalty was chosen by its own inner cross-validation. This penalty was then held fixed while the fusion weight was selected. In each inner fold, the PoE-VAE and probe were refitted on the inner-training patients, as was the standardisation of the latent means. Feature imputation and scaling used the outer-training statistics. No step used the outer test patients.

#### Elastic-net late fusion and concatenation

Both models use elastic-net Cox regression [33], with a mixing parameter of 0.5 and a penalty selected from a path of 30 values by inner five-fold cross-validation. Late fusion fits a separate model to each block using training patients with data for that block, standardises its risk scores on those patients and averages the scores across each patient’s available blocks. Concatenation fits a single model to the concatenated blocks, replacing absent or masked blocks with zeros after standardisation, equivalent to imputing their training means.

#### Neural late fusion

Each block has a network with a single hidden layer of 128 units, trained using the Cox partial likelihood with Breslow handling of ties [25, 34, 35]. Each network is fitted on training patients with data for its block for 100 steps with the same optimiser settings as the PoE-VAE, and the block-specific risk scores are combined as in elastic-net late fusion.

#### Tuned deep models

The PoE-VAE and neural late fusion were also fitted with hyperparameters chosen by inner five-fold cross-validation on the training patients. The candidates, 20 for the PoE-VAE and 12 per block for neural late fusion, varied the number of training steps and the network size and included the fixed configuration (Supplementary Table S2). The candidate with the highest mean inner concordance was refitted on all training patients, with ties resolved towards fewer steps and then the smaller network. Inner folds were formed from the outer-training patients after the outer-training imputation and scaling had been applied.

#### PriorityLasso

We used the R package prioritylasso 0.4.0 [36] with elastic-net penalties and a mixing parameter of 0.5, including penalisation of the first block. Each block’s penalty was selected by minimising five-fold cross-validated deviance, and blocks were ordered by decreasing cross-validated concordance of single-block models. Offset imputation was used for training patients missing a block, whereas missing test blocks were set to zero. An indexing error in the package was corrected (Supplementary Note 2).

#### BlockForest

We used the R package blockForest 0.2.7 [37] with the extremely randomised trees split rule, replacing absent blocks with their training means. We evaluated 50 candidate sets of block weights using 500 trees per set and fitted a final forest of 1,000 trees. The tuning budget and earlier development exposure of every model are listed in Supplementary Table S2.

#### Clinical model

The clinical model was a ridge Cox model [31] with a penalty of 0.1, using age and, where available for the cancer, sex, stage and grade. Variables were chosen by availability without outcomes, and preprocessing was fitted on the training patients (Supplementary Table S1 and Supplementary Note 2).

### 2.4 Outcomes

The endpoint was disease-specific survival, which counts deaths attributed to the cancer, whereas overall survival, the usual endpoint in this field [3–5], also counts deaths from other causes. It was selected after an exploratory comparison of overall, disease-specific and progression-free survival using elastic-net Cox models in KIRC, HNSC, BRCA and LGG, before any model in this study was fitted to the other nine target cancers.

Harrell’s concordance [38] was computed for each test fold, averaged across folds within each repeat and then across repeats. Under natural availability, each test patient was scored using their observed blocks, without artificial masking. A post hoc analysis that masked blocks at test time is described with Supplementary Figure S1.

Uno’s concordance [39] used censoring weights estimated from the training fold and a truncation time set to the 80th percentile of event times for the cancer. The integrated Brier score [40] used the same censoring weights and a grid of 50 equally spaced times between the 10th and 80th percentiles of event times. This grid was shared across models and defined using event times from all patients in the cancer. The integrated Brier score was averaged across test patients within each repeat and then across repeats. Both Uno’s concordance and the integrated Brier score were secondary measures, and integrated Brier scores were also compared with a Kaplan–Meier curve fitted to the training patients [41].

Survival probabilities for the integrated Brier score were obtained from a cross-fitted Cox mapping of each model’s risk score [25, 35, 38] (Supplementary Note 2); BlockForest supplied its own curves, and none were computed for PriorityLasso.

### 2.5 Statistical analysis

For each cancer, we compared models using the difference in mean concordance. We estimated its variance and 95% interval from 2,000 patient-level bootstrap replicates [42]. Patients were sampled with replacement, keeping each patient’s predictions from all five repeats together and using the same resample for both models. Models were not refitted, and predictions, censoring weights and survival mappings remained fixed. A resample was redrawn if any test fold had no comparable patient pair. We then combined the cancer-specific differences in a precision-weighted random-effects model, which allows the underlying difference to vary between cancers. Between-cancer variance was estimated by the DerSimonian–Laird method [43], and pooled intervals by the Hartung–Knapp–Sidik–Jonkman method [44, 45]. The primary comparison was the borrowing model minus the PoE-VAE. Benefit required both a 95% interval entirely above zero and an estimated gain of at least 0.01 in concordance. Molecular model comparisons were grouped into families of two co-primary contrasts (Supplementary Note 3), each using 97.5% intervals to account for the two comparisons by Bonferroni correction. An interval entirely above zero indicated superiority. Otherwise, a lower bound above −0.01 established non-inferiority: a loss of 0.01 or more was excluded. The non-inferiority margin was specified as an analytic tolerance, not a clinically validated threshold. Clinical-model comparisons were interpreted by whether the interval lay above or below zero. An interval including zero did not establish equivalence. Uno’s concordance used the same interval levels as Harrell’s concordance; intervals for the integrated Brier score and individual cancers were unadjusted.

External comparisons tested the PoE-VAE against elastic-net late fusion, neural late fusion and PriorityLasso. These three prespecified contrasts used 98.33% intervals, applying Bonferroni correction for three comparisons. The 0.02 gain used for comparison with published studies [10, 11] served only as a reference for comparison with the literature; it was not a decision threshold in this study.

The two tuned deep models were each compared with elastic-net late fusion using 97.5% intervals and the superiority and non-inferiority rules above, including the 0.01 margin. For VAECox, the primary comparison was the pretrained minus the randomly initialised model, with the target cancer excluded from pretraining (Section 2.8). Benefit required a 95% interval entirely above zero and an estimated gain of at least 0.01. Other protocol versions and differences between consecutive versions were descriptive.

Exploratory follow-ups repeated the analysis in a ten-cancer overall-survival setting closer to the original study and, in both settings, evaluated each network at the fine-tuning step chosen using inner validation data (Section 2.8).

Analyses were specified in stages. Here, “prespecified” means written down before that stage was run; later specifications were informed by earlier results from the same patients (Supplementary Note 3). Post hoc analyses included an equally weighted mean across the 13 cancers, with a percentile interval obtained by averaging their bootstrap draws while treating the cancers as fixed. We also calculated modified Hartung–Knapp intervals [46], constraining the scale factor to be at least one. Two further post hoc analyses used stored predictions to examine unequal test-step selection in VAECox (choosing the best test step for only one network) and variation in model comparisons between single five-fold runs. Both were specified after their point estimates had been inspected. Supplementary Note 3 records the analysis chronology; other post hoc analyses are labelled where reported.

### 2.6 Site-held-out folds

Tissue source sites were identified from the second field of the TCGA barcode and assigned to folds following the principle of preserved-site cross-validation [47]. Sites were ordered by decreasing event count, with ties broken using a seeded random ordering, and each was assigned in full to the fold with the fewest events so far, favouring the fold with fewer patients when event counts were equal. Five-fold partitions were redrawn up to 20 times if any fold contained fewer than five events, and models were refitted on these folds using the same training procedures.

### 2.7 External cohorts

CPTAC-3 matrices and clinical data [48] were obtained from the GDC Xena hub [2, 21]. Six cohorts matched a TCGA target and had at least 20 deaths. The external outcome was overall survival, whereas models were trained for disease-specific survival. Recorded sex was checked against RNA expression of XIST and RPS4Y1 and against X-chromosome methylation; every cohort passed the check, and no patient was excluded on it (Supplementary Table S1). Models were trained once per seed on all eligible TCGA patients of the cancer.

We evaluated three preprocessing pipelines. The primary cohort-adapted pipeline used external-cohort data to select available features and to impute and scale external measurements. The prespecified scaling sensitivity kept the adapted features but used TCGA preprocessing statistics. The exploratory fully frozen pipeline used both the TCGA feature set and TCGA preprocessing statistics.

For the primary pipeline, a feature was retained if at least 80% of patients with the corresponding external assay had a finite measurement and its standard deviation after external-cohort median imputation exceeded 10^−8^. Models were trained on the retained features. Within each observed external block, sporadically missing values were imputed using the medians of eligible CPTAC-3 patients with that block. Features were then centred and scaled using that cohort’s post-imputation means and standard deviations.

The scaling sensitivity used the same adapted feature set and fitted models. Only external preprocessing changed: missing values were imputed with TCGA training medians, and scaling used TCGA training means and standard deviations.

In all external configurations, a patient without a modality sample retained an absent block rather than receiving an imputed assay. The PoE-VAE and the pretrained encoders used only observed blocks; elastic-net and neural late fusion averaged risks over observed blocks; PriorityLasso set missing test blocks to zero after standardisation. A block absent from the corresponding TCGA training cohort remained unavailable for external prediction, including microRNA in GBM.

The fully frozen pipeline retained the full TCGA feature set. Within observed external blocks, missing values and features absent from the external platform were imputed with TCGA training medians. Scaling used TCGA training means and standard deviations. This follow-up was exploratory (Supplementary Note 3), and comparisons between preprocessing pipelines were descriptive.

### 2.8 Published borrowing method (VAECox)

We reimplemented VAECox [10] from its public code (github.com/dmis-lab/VAECox). It tests borrowing with cohort-scale target training using RNA data available to us, and its reported gain of 0.020 is one of our two reference values. VAECox first pretrains a variational autoencoder on pooled RNA profiles from multiple cancers, then fine-tunes its encoder for survival prediction. As in the original study, the reference is the same network with a randomly initialised encoder.

We followed the published architecture: an autoencoder with one hidden layer of 4,096 units and a 128-dimensional mean, followed by a Cox-nnet head with one hidden layer of 12 units. We also followed the pretraining objective and schedule (501 full-batch Adam steps) and the 21 fine-tuning steps under the Cox partial likelihood. Inner five-fold cross-validation selected one of six combinations of learning rate and weight decay. Each candidate was scored by its highest mean validation concordance across fine-tuning steps.

Two choices in the published code were of particular interest. Pretraining draws a random 80% sample of the pooled patients from 20 cancers, independently of the survival splits, so most target-cancer test patients enter pretraining. The code also records test concordance after every fine-tuning step and reports the maximum. To examine these choices and the use of test patients in scaling, we ran four versions using each target’s RNA feature set, disease-specific survival and our partitions:

1. Published-style procedure: pretrain on an 80% sample of the pooled patients from all 20 cohorts, including target-cancer test patients; scale using all target patients; and evaluate at the best test step.
2. Last-step evaluation: use the same fitted models as in version (1), but evaluate at the last fine-tuning step.
3. Test-patient exclusion: retain last-step evaluation, exclude the partition’s test patients from pretraining and estimate scaling from training patients.
4. Target-excluded protocol: also exclude every other patient of the target cancer from pretraining, retaining training-patient scaling and last-step evaluation.

Within each cancer and seed, the pretrained and randomly initialised encoders started from the same weights. All versions were scored on the same test patients with RNA data. Supplementary Note 2 gives implementation details and deviations from the original study.

Versions (2) to (4) evaluate the last fine-tuning step, although the hyperparameter search scores candidates at their best mean validation step. We therefore added a validation-selected evaluation of the target-excluded protocol. For the chosen candidate, we used the step with the highest mean inner validation concordance, taking the first step if several tied. This step was selected separately for each network and partition, without using test patients.

Our features, endpoint and pretraining cohorts differed from those in the original study. We therefore repeated all four versions and the validation-selected evaluation in an exploratory ten-cancer overall-survival setting closer to that study. We used VAECox’s ten target cancers and 20 pretraining cancers, adding RNA from THCA, SKCM, PRAD and LAML.

The shared feature set contained 18,858 protein-coding genes, selected without outcomes. Each gene was present in all 20 cohorts, had at most 20% missing values and had positive expression in at least 10% of the pooled samples. All versions used this feature set and new five repeats of five-fold cross-validation among patients with RNA data and an overall-survival label (Supplementary Note 2).

### 2.9 Software

Experiments were performed on the computational facility of the National Computational Infrastructure (NCI) through the UNSW HPC Scheme (DOI: 10.26190/PMN5-7J50). Analyses used Python 3.10, PyTorch 2.7 [49], scikit-survival 0.25 [50] and R 4.4.2.

## 3 Results

### 3.1 Study design

We included the 13 TCGA cancer cohorts with at least 75 disease-specific deaths among patients with molecular data (Table 1). Features were selected in the other 19 cohorts (4,000 RNA, 528–576 microRNA and 4,000 methylation features per target). In GBM and OV one block was not used because fewer than 50 patients had it (microRNA and methylation, respectively). Of the 5,761 patients, 4,297 (75%) had all three blocks; 206 (4%) lacked RNA, 323 (6%) lacked microRNA and 1,245 (22%) lacked methylation data (Supplementary Table S1).

**Table 1.**
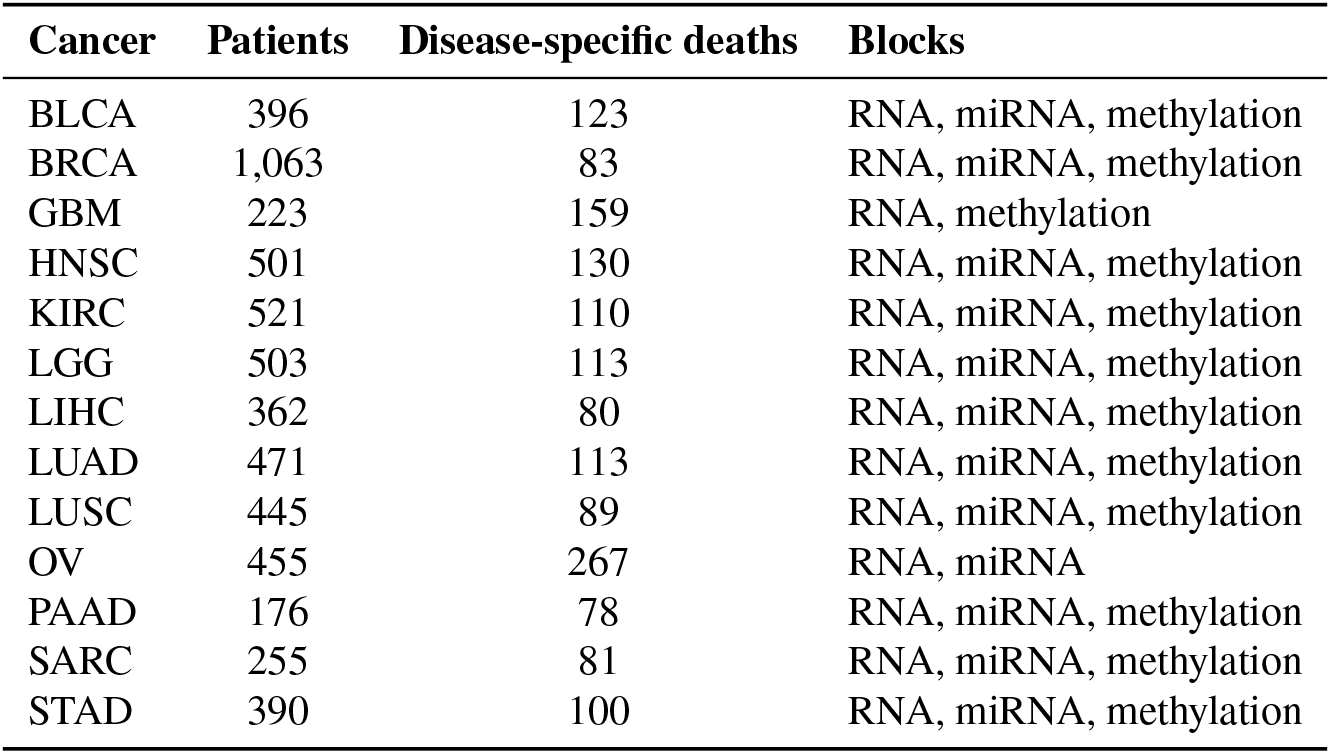
Target cancers. Patients and deaths are those eligible for the disease-specific survival analysis. BLCA, bladder urothelial carcinoma; BRCA, breast invasive carcinoma; GBM, glioblastoma multiforme; HNSC, head and neck squamous cell carcinoma; KIRC, kidney renal clear cell carcinoma; LGG, brain lower grade glioma; LIHC, liver hepatocellular carcinoma; LUAD, lung adenocarcinoma; LUSC, lung squamous cell carcinoma; OV, ovarian serous cystadenocarcinoma; PAAD, pancreatic adenocarcinoma; SARC, sarcoma; STAD, stomach adenocarcinoma. miRNA, microRNA.

| **Cancer** | **Patients** | **Disease-specific deaths** | **Blocks** |
|---|---|---|---|
| BLCA | 396 | 123 | RNA, miRNA, methylation |
| BRCA | 1,063 | 83 | RNA, miRNA, methylation |
| GBM | 223 | 159 | RNA, methylation |
| HNSC | 501 | 130 | RNA, miRNA, methylation |
| KIRC | 521 | 110 | RNA, miRNA, methylation |
| LGG | 503 | 113 | RNA, miRNA, methylation |
| LIHC | 362 | 80 | RNA, miRNA, methylation |
| LUAD | 471 | 113 | RNA, miRNA, methylation |
| LUSC | 445 | 89 | RNA, miRNA, methylation |
| OV | 455 | 267 | RNA, miRNA |
| PAAD | 176 | 78 | RNA, miRNA, methylation |
| SARC | 255 | 81 | RNA, miRNA, methylation |
| STAD | 390 | 100 | RNA, miRNA, methylation |

### 3.2 Estimated borrowing gains are small overall and depend on cancer weighting

In the prespecified analysis, the change in concordance from borrowing, the borrowing model minus the PoE-VAE (Section 2.3), was +0.002 (95% interval −0.002 to +0.006; Figure 2a; per-cancer values in Supplementary Table S3 and pooled values in Supplementary Table S4). The between-cancer variance was estimated at zero, and the interval, which conditions on the fitted models, excluded a gain of 0.01. All cancer-specific intervals included zero, with the largest point estimates in LUAD (+0.033, −0.013 to +0.080) and HNSC (+0.025, −0.006 to +0.058).

**Figure 2.**
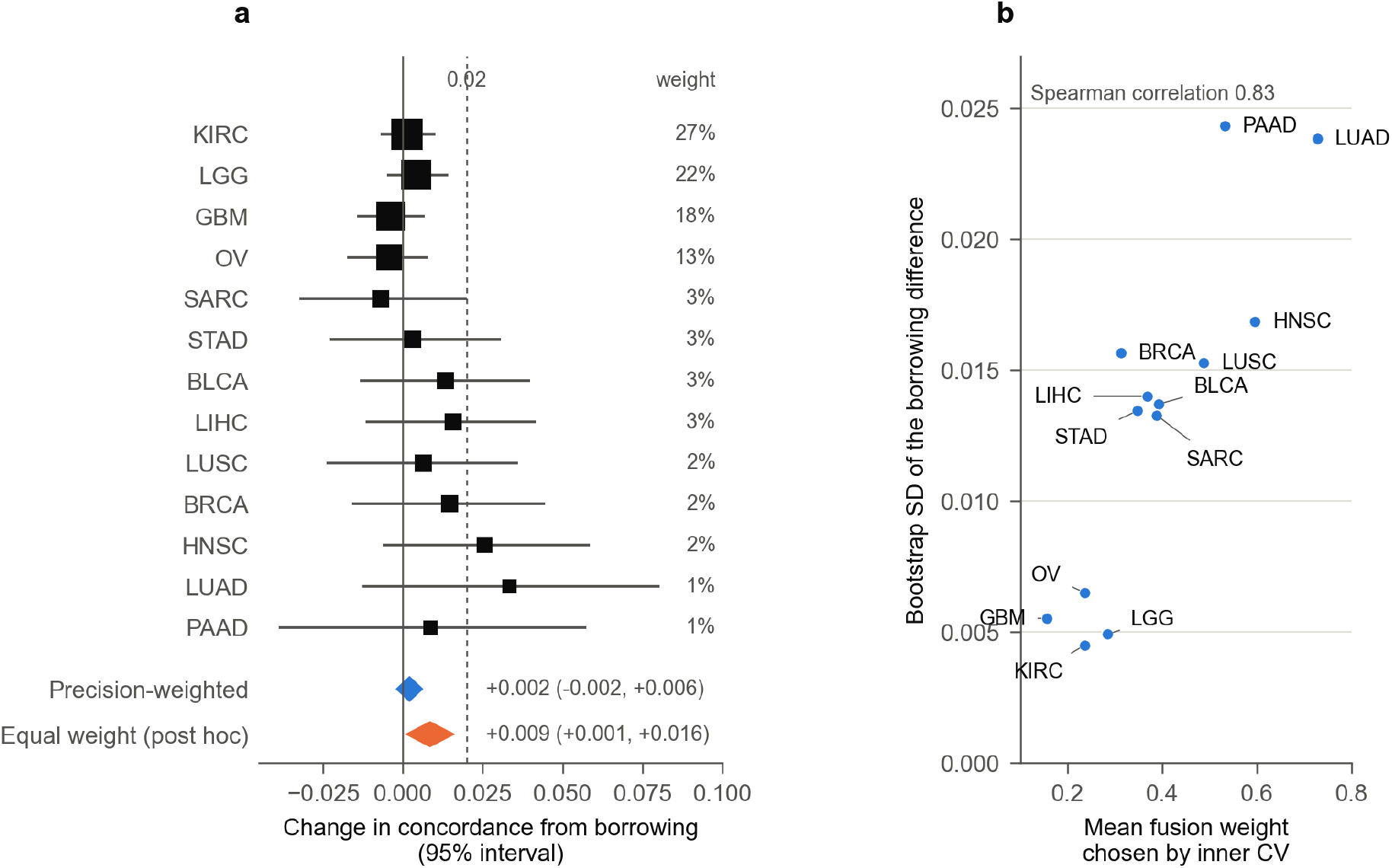
Borrowing across cancers. (a) Change in concordance from borrowing (the borrowing model minus the PoE-VAE, disease-specific survival, natural availability) in each cancer, with 95% paired bootstrap intervals. Cancers are ordered by their weight in the random-effects analysis (right-hand column), and marker area follows that weight. Diamonds show the prespecified precision-weighted estimate (Hartung-Knapp-Sidik-Jonkman interval) and the post hoc estimate with equal weight per cancer (percentile interval). The dashed line marks 0.02, the smaller of the two published gains used for comparison [10, 11]. (b) The bootstrap standard deviation (SD) of the borrowing difference against the mean fusion weight chosen by inner cross-validation (CV). Cancers in which little weight was given to the borrowed score have the smallest standard deviations and therefore the largest weights in (a).

The random-effects summary weights each cancer by the estimated uncertainty of its difference and the between-cancer heterogeneity, and four cancers (KIRC, LGG, GBM and OV) received 80% of the weight. These were also the cancers in which inner cross-validation selected the smallest fusion weights, averaging 0.16–0.28, so the fused model was close to the reference and the difference between them had a small variance. Across the 13 cancers, the Spearman rank correlation between mean fusion weight and the bootstrap standard deviation of the difference was 0.83 (Figure 2b). The random-effects summary was therefore dominated by cancers in which little borrowing occurred. The arithmetic mean across the 13 cancers, a post hoc summary whose interval treats these cancers as fixed, was +0.009 (+0.001 to +0.016). Equal weighting gives more influence to the cancers in which the model made more use of the pretrained score.

The estimate was +0.0005 (95% interval −0.0048 to +0.0059) after excluding the three cancers for which the curators caution against disease-specific survival (BRCA, LGG and LIHC [6]; post hoc; Supplementary Table S4) and +0.0005 (−0.0049 to +0.0059) with Uno’s concordance.

### 3.3 A reimplementation of VAECox shows no average pretraining gain under matched evaluation

Pretraining showed no average gain over random initialisation when both networks were evaluated alike (Supplementary Table S5). In the 13-cancer analysis, the published-style procedure gave a concordance difference of −0.004 (95% interval −0.010 to +0.001). We also applied the original study’s summary: an average over VAECox’s ten cancers weighted by patient count. This gave −0.0055 (concordance 0.7126 for pretraining and 0.7181 for random initialisation), compared with the reported gain of +0.020 [10].

Under the target-excluded protocol, the difference was −0.004 (−0.011 to +0.003) when both networks were evaluated at the last step. It was −0.001 (−0.012 to +0.010) when each network was evaluated at its validation-selected step. Both intervals excluded a gain of 0.02; the last-step interval also excluded a gain of 0.01. Differences within individual cancers were larger in both directions at the validation-selected step (Supplementary Table S5).

Between successive versions of the procedure (Section 2.8), the difference changed by about 0.003 or less, with every interval including zero (Supplementary Table S5).

Pretraining also failed to improve average concordance in the exploratory ten-cancer overall-survival setting (Section 2.8; Supplementary Table S5). The published-style procedure gave a difference of −0.006 (−0.011 to −0.001). Weighting these ten cancers by patient count, as in the original study, gave −0.0063 (concordance 0.6713 for pretraining and 0.6776 for random initialisation). Under the target-excluded protocol, the differences were −0.009 (−0.016 to −0.003) at the last step and −0.004 (−0.008 to −0.0002) at the validation-selected step.

Moving from the target-excluded protocol towards the published-style procedure shifted the estimate towards pretraining by 0.001 to 0.003 at each successive change. None of these shifts was distinguishable from zero. Together, the three changes shifted the estimate by about 0.004, well below the reported gain of 0.020.

### 3.4 Evaluation choices change differences between models and absolute concordance

Selecting a single fusion weight per cancer using the test results increased the equal-weighted borrowing estimate by 0.008, to 0.017 (Table 2). Under the prespecified random-effects summary, the estimate increased from 0.002 to 0.007 (+0.003 to +0.011). Intervals for these test-selected estimates condition on the selection and do not account for the search over weights [51].

**Table 2.** Size of evaluation effects. Each row gives the evaluation choice, what it changes, the quantity it affects and an estimate, identified in each row as a change or as the resulting value. Changes given without an interval are differences between two reported means. Unless stated, sizes are means across cancers with equal weight and 95% intervals; these summaries and the rows marked “post hoc” were added after the results were available. VAECox rows use our reimplementation on the 13 cancers, and “ten-cancer setting” is the ten-cancer overall-survival analysis (Section 2.8). Intervals for test-selected quantities condition on the selection. The unequal-selection rows are a constructed experiment; the published VAECox procedure evaluates every model at its best test step.

| Evaluation choice | What changes | Quantity affected | Estimate |
|---|---|---|---|
| Fusion weight chosen per cancer on the test splits | The selection procedure | Difference between models (borrowing gain) | Change +0.008 (+0.005, +0.011) from the cross-validated weight, positive in 12 of 13 cancers (post hoc); resulting gain +0.017 (+0.009, +0.025), against +0.009 |
| Pretrained VAECox network evaluated at its best test step, the other at its last step (post hoc) | Whether the two networks receive the same treatment | Difference between models (pretrained minus randomly initialised) | Change +0.049, from −0.004 with both at the last step to +0.045 (+0.035, +0.054); ten-cancer setting: change +0.027, from −0.005 to +0.022 (+0.014, +0.030). Intervals are for the resulting difference |
| Randomly initialised network evaluated at its best test step, the other at its last step (post hoc) | Whether the two networks receive the same treatment | Difference between models (pretrained minus randomly initialised) | Change −0.047, to −0.051 (−0.059, −0.042); ten-cancer setting: change −0.027, to −0.032 (−0.039, −0.025). Intervals are for the resulting difference |
| Both VAECox networks evaluated at their best test step | The selection procedure, applied to both networks | Absolute concordance | Change +0.047 and +0.049 for the two networks (+0.027 in the ten-cancer setting); their difference changed by 0.002 |
| One five-fold run reported instead of five (post hoc) | The particular partitions and fitted models | Difference between models | SD 0.012 for the borrowing gain and 0.012 to 0.022 for six model comparisons; 11% and 12% to 42% of runs at least 0.02 from the five-run mean; SD 0.009 to 0.020 and 0% to 23% of runs for four VAECox comparisons |
| Arithmetic mean across cancers instead of the random-effects summary (post hoc) | The cross-cancer summary | Difference between models (borrowing gain) | Mean +0.009 (+0.001, +0.016), against +0.002 (−0.002, +0.006) for the random-effects summary |
| Target cancer’s patients added, unlabelled, to pretraining | The pretraining data | Difference between models (borrowing gain) | Change 0.000 (−0.003, +0.003) |
| Whole tissue source sites held out (Section 3.5) | The evaluation population and the partition design | Absolute concordance | Change −0.023 (−0.040, −0.006) for the PoE-VAE and −0.016 (−0.029, −0.002) for elastic-net late fusion (random-effects summary, 97.5% intervals) |

Choosing the best test step for both VAECox networks raised absolute concordance but had little effect on their difference (Table 2). This is the selection rule used for every model in the published code. In the 13-cancer analysis, absolute concordance increased by 0.047 to 0.049 on average and by up to 0.09 in individual cancers. The average increase was 0.027 in the ten-cancer overall-survival setting. In both settings, the difference between networks changed by about 0.002 or less. Concordance at the best test step includes the effect of searching over steps and is therefore not directly comparable with concordance at a fixed or validation-selected step.

Choosing the best test step for only one network had a much larger effect on the difference. We examined this in a constructed post hoc experiment using the same fitted networks. Unlike the published VAECox procedure, this experiment applied unequal selection: the pretrained network was evaluated at its best test step, and the randomly initialised network at its last step.

In the 13-cancer analysis, the difference (pretrained minus randomly initialised) changed from −0.004 with both networks at the last step to +0.045 (95% interval +0.035 to +0.054). This was a shift of +0.049, and the resulting difference was positive in 12 of 13 cancers. In the ten-cancer overall-survival setting, the difference changed from −0.005 to +0.022 (+0.014 to +0.030), a shift of +0.027. Selecting the best test step only for the randomly initialised network instead gave differences of −0.051 and −0.032, respectively. The apparent advantage therefore followed which network received test-step selection (Supplementary Table S5).

Including the target cancer’s patients in pretraining without their outcomes had no detectable effect (Table 2).

Single-run estimates varied substantially in the post hoc analysis. For the borrowing gain, estimates from single five-fold runs had a standard deviation of 0.012 around each cancer’s five-run mean. Of the 65 runs, 11% differed from that mean by at least 0.02. Estimates from single train–test partitions varied about three times as much (Supplementary Table S6).

Variation was at least as large across six comparisons among the PoE-VAE, elastic-net models, neural late fusion and PriorityLasso. Their single-run standard deviations ranged from 0.012 to 0.022, and 12% to 42% of runs differed from the corresponding five-run mean by at least 0.02 (Figure 3; Supplementary Table S6). Across four VAECox comparisons, the corresponding ranges were 0.009 to 0.020 and 0% to 23%. These estimates combine variation in the partitions with randomness in model fitting.

**Figure 3.**
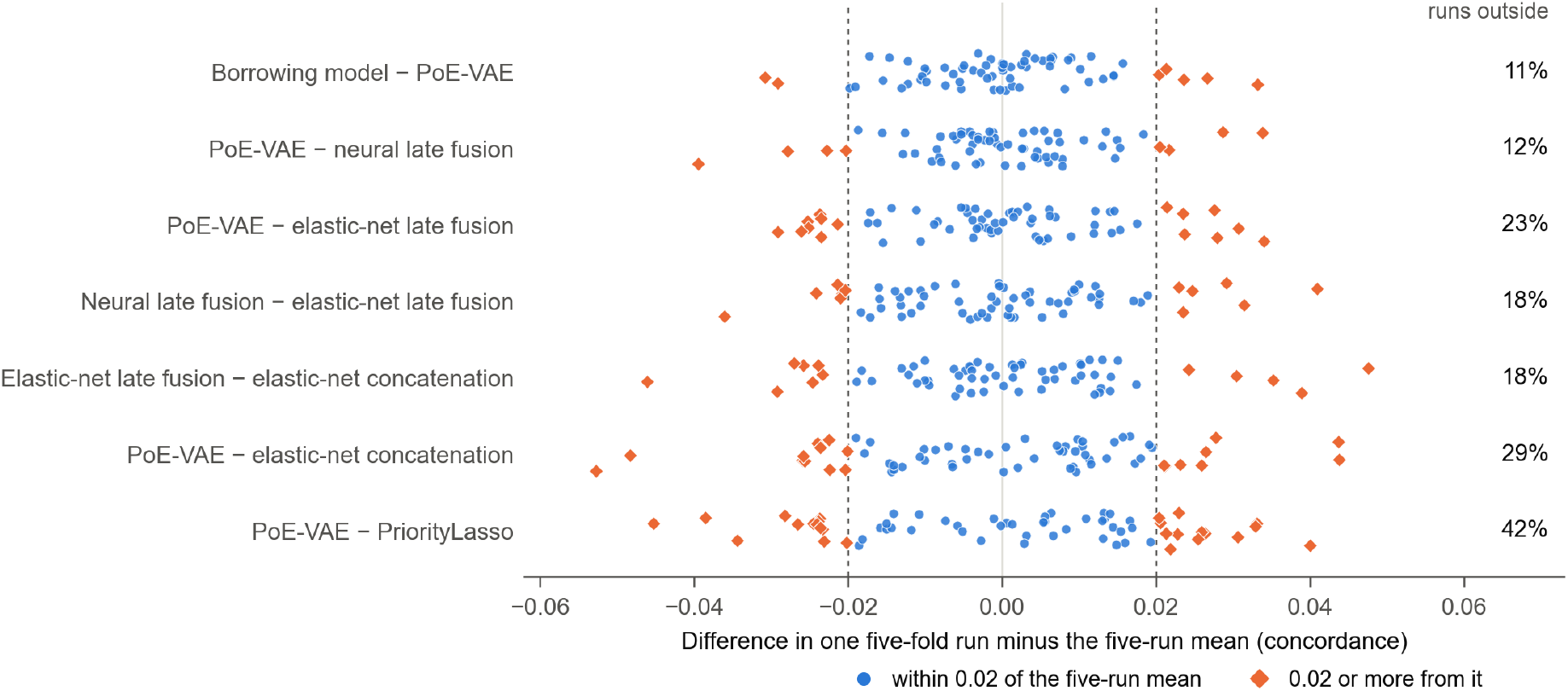
Variation between single five-fold runs (post hoc). Each point is one five-fold run of one cancer (five runs in each of 13 cancers, 65 per row): the difference in concordance between the two models named, under natural availability, minus the mean of that cancer’s five runs. Dashed lines mark deviations of 0.02. Diamonds are runs at least 0.02 from their cancer’s mean, and their share is given on the right. Deviations are measured against the five-run mean, not a known true value; points are spread vertically for legibility. The four VAECox comparisons are reported in Supplementary Table S6.

### 3.5 Concordance is lower on unseen tissue source sites

TCGA samples were collected at many sites, and random folds place patients from the same site in both training and test sets [47]. We rebuilt the folds so that each site fell wholly within one fold, balancing events across folds, and refitted the PoE-VAE and elastic-net late fusion.

Concordance was lower by 0.023 for the PoE-VAE (97.5% interval −0.040 to −0.006) and by 0.016 for elastic-net late fusion (−0.029 to −0.002; Figure 4; Supplementary Table S7). The decrease was largest in LIHC (0.107 and 0.083) and KIRC (0.053 and 0.061) and near zero in BRCA, LUAD, LUSC and STAD. The difference between their decreases was −0.007 (95% interval −0.015 to +0.001); the comparison of the two models on site-held-out folds is given in Supplementary Table S4.

**Figure 4.**
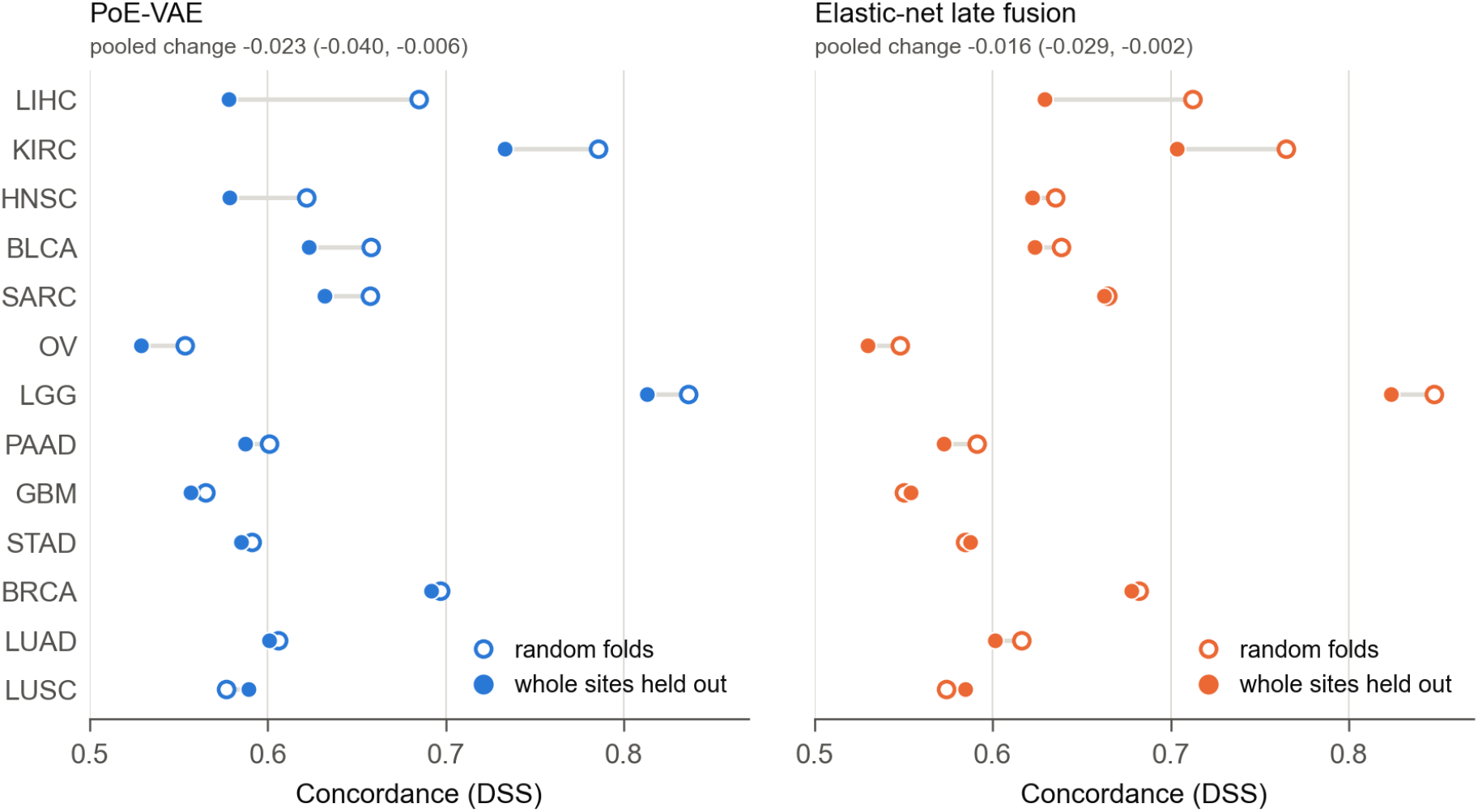
Site-held-out folds. Harrell’s concordance for disease-specific survival per cancer with random folds (open markers) and with whole tissue source sites held out (filled markers), for the PoE-VAE and elastic-net late fusion. Lines connect point estimates and are not intervals. Cancers are ordered by the size of the change for the PoE-VAE. Pooled changes (site-held-out minus random folds) are given with 97.5% intervals. The five seeds share one site partition in KIRC, two in STAD, three in OV and four in LGG.

### 3.6 The reference model is non-inferior to PriorityLasso and BlockForest

To check that the PoE-VAE is a competitive reference model for the borrowing analysis, we compared it with five other models using the same features and splits (Section 2.3; Table 3; per-cancer values in Supplementary Table S3). PriorityLasso and BlockForest are the methods ranked highest by the published benchmarks [3–5].

**Table 3.** Competitiveness of the PoE-VAE reference model. In the first five rows each entry is the fixed PoE-VAE minus the comparator, pooled over 13 cancers under natural availability. Harrell’s concordance is primary. Concordance intervals are 97.5%, assigned within separate families of two contrasts (the PoE-VAE against elastic-net late fusion and concatenation; the PoE-VAE against PriorityLasso and neural late fusion; the PoE-VAE and elastic-net late fusion, the latter not shown, against BlockForest; and the two tuned models against elastic-net late fusion in the last two rows), not jointly over all rows. The integrated Brier score (IBS) is secondary, with nominal 95% intervals; a negative IBS difference favours the PoE-VAE. The IBS contrast with elastic-net concatenation was added post hoc.

| **Comparator or contrast** | **Difference in Harrell’s concordance** | **Difference in Uno’s concordance** | **Difference in IBS** |
|---|---|---|---|
| Elastic-net late fusion | +0.001 (−0.009, +0.012) | +0.005 (−0.007, +0.016) | −0.0003 (−0.0010, +0.0004) |
| Neural late fusion | −0.005 (−0.016, +0.005) | −0.005 (−0.015, +0.005) | +0.0014 (+0.0002, +0.0025) |
| PriorityLasso | +0.014 (−0.009, +0.038) | +0.017 (−0.003, +0.038) | not computed |
| BlockForest | +0.008 (−0.006, +0.022) | +0.015 (+0.0001, +0.029) | −0.0015 (−0.0028, −0.0003) |
| Elastic-net concatenation | +0.018 (+0.003, +0.033) | +0.021 (+0.006, +0.037) | −0.0013 (−0.0026, −0.0001) |
| Tuned PoE-VAE minus elastic-net late fusion | +0.006 (−0.008, +0.019) | not computed | not computed |
| Tuned neural late fusion minus elastic-net late fusion | +0.012 (+0.001, +0.022) | not computed | not computed |

The PoE-VAE achieved higher concordance than elastic-net concatenation; under the prespecified random-effects summary, the other four comparisons were inconclusive for superiority in either direction. With equal weight per cancer, a post hoc summary, the comparison with PriorityLasso favoured the PoE-VAE (+0.013, 97.5% interval +0.0005 to +0.026; Supplementary Table S4). Using the prespecified margin of 0.01, the PoE-VAE was non-inferior to elastic-net late fusion, PriorityLasso and BlockForest.

Non-inferiority to neural late fusion was not established (−0.005, 97.5% interval −0.016 to +0.005). These comparisons support the PoE-VAE as a reference for the borrowing analysis without identifying it as the best-performing model.

With hyperparameters chosen by inner cross-validation (Section 2.3), tuned neural late fusion exceeded elastic-net late fusion by 0.012 (97.5% interval +0.001 to +0.022), with lower bounds above zero also under equal cancer weighting and the modified Hartung–Knapp interval. The tuned PoE-VAE differed from elastic-net late fusion by +0.006 (97.5% interval −0.008 to +0.019) and from its fixed configuration by +0.002 (95% interval −0.006 to +0.009). It was 0.007 below tuned neural late fusion under the random-effects summary (95% interval −0.013 to −0.0001); with equal cancer weighting the interval included zero. Inner cross-validation most often selected 10 training steps, the shortest candidate, and rarely the fixed configuration (Supplementary Table S2).

For context, a ridge Cox model using up to four clinical variables (age, sex, stage and grade) had a higher point estimate of concordance than the fixed PoE-VAE in 9 of the 13 cancers; the pooled difference, PoE-VAE minus clinical model, was −0.020 (97.5% interval −0.055 to +0.016), consistent with Herrmann et al. [3] (Supplementary Tables S3 and S4).

### 3.7 In CPTAC-3, concordance is 0.58–0.60 and the prespecified model comparisons are inconclusive

Models trained for disease-specific survival on all eligible TCGA patients of each cancer were evaluated in the corresponding CPTAC-3 cohort, 965 patients and 431 deaths across six cancers, for overall survival because cause of death was incompletely recorded. In the primary cohort-adapted pipeline, features were filtered and scaled using each external cohort’s own data, without outcomes (Section 2.7); comparisons with internal TCGA concordance are descriptive because the endpoint, feature availability and training size differ.

Pooled concordance ranged from 0.58 to 0.60 across models, with all intervals excluding 0.5 (Supplementary Table S7). Concordance in LUAD, LUSC, GBM and PAAD was close to the cross-validated estimates in TCGA, whereas in KIRC it was 0.08–0.12 lower for every model. KIRC was also one of the two cancers with the largest decreases under site-held-out evaluation. The three prespecified comparisons were inconclusive, with 98.33% intervals for the PoE-VAE minus the comparator of −0.033 to +0.015 for elastic-net late fusion, −0.036 to +0.017 for neural late fusion and −0.049 to +0.060 for PriorityLasso (Supplementary Table S7). The estimated gain from borrowing was +0.007 (−0.001 to +0.015).

The three prespecified comparisons remained inconclusive with TCGA scaling statistics and in the exploratory fully frozen pipeline, where the borrowing gain was +0.008 (95% interval +0.0001 to +0.015; Supplementary Figure S2 and Supplementary Table S7).

## 4 Discussion

Evaluation choices changed estimated gains by amounts comparable to the reported concordance gains of 0.020 and 0.045, although those published gains come from different tasks. In the constructed post hoc experiment, choosing the best test step for only one VAECox network shifted the mean difference by 0.027 to 0.049 towards that network. Choosing the fusion weight on test splits raised the equal-weight borrowing gain by 0.008. A single five-fold run could also show or hide a gain of this size: across seven multi-omics comparisons, 11% to 42% of runs differed from their cancer’s five-run mean by at least 0.02.

Choosing the best test step for both VAECox networks raised absolute concordance by 0.03 to 0.05 but changed their difference by about 0.002 or less. This cancellation was observed for these networks and may not hold when models have different learning curves. Under the reference design, our borrowing configuration and the VAECox reimplementation showed little or no average gain. The estimate for our borrowing model also depended on how cancers were weighted.

Holding out tissue source sites is a different kind of choice: it changes the evaluation population and asks how a model performs at sites not represented in training. Concordance was about 0.02 lower there, and the comparison of the two models’ decreases was inconclusive. Site signatures in TCGA histology [47] and unwanted technical variation in TCGA RNA-seq [52] have been reported, but our analysis does not identify the cause of the decrease.

Previous work has shown that evaluation choices can change benchmark conclusions. Using multi-omics survival prediction as an example, Nießl et al. [12] showed that rankings depend on the datasets, performance measure, handling of missing performance values and aggregation method. SurvBoard standardises evaluation across datasets [5].

Here, we distinguish the effect of test-set selection on the difference between models from its effect on absolute concordance. We measure both effects in our borrowing configuration and a reimplementation of a published method, and compare their size with reported gains. We also show why precision weighting can favour cancers in which a model’s learned fusion weight produces little change from its reference. For the pipelines tested, our model comparison agrees with the three benchmarks [3–5]: the prespecified summary did not establish that the PoE-VAE outperformed block-wise references (Section 3.6).

The borrowing findings are limited to the two implementations evaluated. For our configuration, both pooled intervals excluded an average gain of 0.02; the equal-weight lower bound was just above zero, and the cancer-specific intervals allowed gains of several hundredths in individual cancers. The VAECox reimplementation, which fine-tunes a pretrained encoder, also showed no gain at the last or the validation-selected step, and in the ten-cancer overall-survival setting pretraining slightly lowered concordance. It differs from the original experiment in expression processing, resampling and the reconstructed randomly initialised reference (Supplementary Note 2). Gross et al. [20] likewise found no benefit from autoencoder pretraining with the downstream cohorts excluded. Several studies reporting an overall benefit also described settings in which transfer did not help [7, 10, 16]. Sabbatini et al. [8], whose design is closest to ours (RNA, the target tumour excluded from pretraining, then fine-tuning), reported a benefit of transfer in 24 of 27 tumour types for disease-free survival. Their released notebooks evaluate candidate configurations on held-out splits without a separate validation set (Supplementary Note 1).

The borrowing estimate was +0.002 with random-effects pooling and +0.009 with equal cancer weighting. When a learned combination can fall back on its reference model, as with a fusion weight close to zero, the cancers in which this occurs yield differences with small variances and receive most of the inverse-variance weight. Reporting both summaries makes this dependence visible and lets readers judge which one answers their cross-cancer question.

These experiments quantify evaluation effects in specified pipelines; they do not explain any particular published result. For VAECox, the departures of the published-style procedure from the target-excluded protocol moved the difference by less than 0.005, and the published code treats all its models alike. What the experiments show is that a reported gain of a few hundredths cannot be interpreted on its own, and three things need to be reported with it. The first is how each model was selected: hyperparameters and training length chosen without the test data, with comparable selection opportunities and a documented budget for every model compared [32, 51]. The second is the uncertainty of the paired difference: intervals for differences on the same test patients and the variation across repeated partitions. The third is the evaluation population and the summary: which patients or sites were held out and, when pooling across datasets, the arithmetic mean alongside a precision-weighted summary. These details let readers judge whether a reported gain could reflect evaluation effects of the size measured here.

### Limitations

The data limit the scope of the findings. All internal analyses use TCGA, where disease-specific survival is approximated from tumour status in most cancers. The curators caution against this endpoint in three of the 13 cancers (BRCA, LGG and LIHC) [6]. Our sensitivity analysis excluding them covers only the borrowing contrast and six model and feature-selection contrasts (Supplementary Table S4).

The external cohort shares the GDC processing pipeline [21]. It was evaluated for overall survival rather than the disease-specific endpoint used in training, and the primary analysis adapted its feature set and scaling using cohort data without outcomes. Site-held-out folds change case mix and training-set size as well as site representation, so the decrease cannot be attributed solely to site-specific signal. The site constraint also caused partitions to repeat across seeds in four cancers (Figure 4).

The uncertainty estimates are conditional on the data and fits used here. Bootstrap intervals do not include variation from refitting models on new patients, so their exclusion of gains of 0.01 or 0.02 applies to these fits (Supplementary Note 2). Equal-weighted intervals treat the 13 cancers as fixed, and site-held-out intervals condition on the observed sites.

There are also limitations to model selection and analysis planning. Preprocessing and the probe penalty were fitted on each outer-training set and reused in inner cross-validation. The inner scores therefore do not represent a pipeline refitted entirely within each inner fold, although no outer test data were used. The endpoint was selected after an exploratory comparison in four cancers, and the PoE-VAE was developed on KIRC and HNSC. Later analysis stages used the same patients and were specified with knowledge of earlier results. The unequal-selection experiment and the single-run analysis of model comparisons were specified after their point estimates had been inspected (Supplementary Note 3). Interval adjustments cover the named contrasts within each stage only; they provide no study-wide guarantee.

The findings apply to the configurations tested. We compared one PoE-VAE architecture and one neural late-fusion architecture, each with fixed and tuned hyperparameters. These architectures differ in likelihood, baseline hazard and training on subsets of blocks, so the comparison cannot isolate the contribution of the generative encoder or product-of-experts fusion. BlockForest also searched fewer weight sets than its default (Supplementary Table S2).

We did not vary missingness in the training data, and late fusion and concatenation handle incomplete training patients differently. The supplementary masking analysis used fixed fitted models and complete-case patients. It therefore does not reproduce the population with naturally missing blocks, for whom missingness may be informative [53].

The borrowing analysis fused a frozen pretrained representation with the fixed reference model; it did not test borrowing with the tuned reference. VAECox was reimplemented using our features and endpoint, with training length selected within the published 21 fine-tuning steps. It was not rerun on its original data, and other published borrowing methods were not rerun. Finally, the molecular models used only omics data. Because we did not combine clinical variables with omics, the study does not assess the prognostic value of omics beyond clinical variables.

---

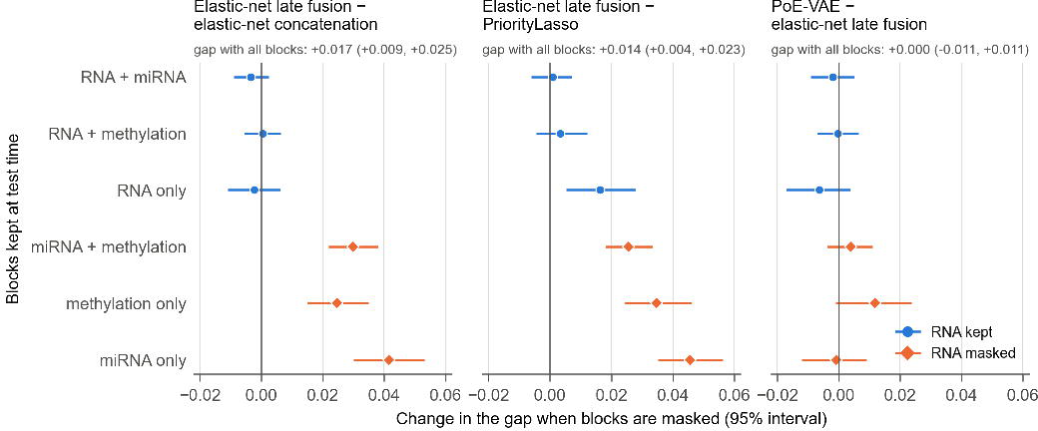

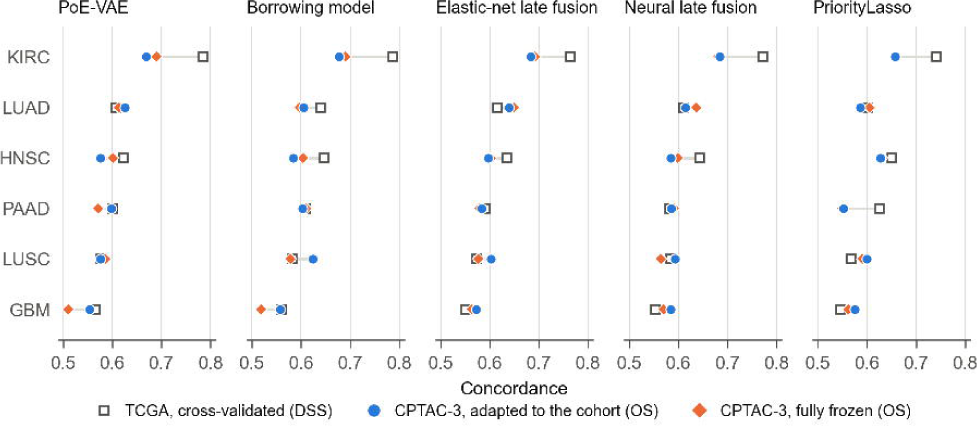

