## Supplementary Material for "Evaluation effects can rival reported concordance gains in cross-cancer survival prediction"

Tables: S1, cohorts, block availability and fold sizes; S2, model selection and tuning budget; S3, concordance, borrowing gains and integrated Brier scores per cancer; S4, pooled paired contrasts and sensitivity analyses; S5, VAECox reimplementation; S6, variation between single runs; S7, site-held-out and external evaluation. Figures: S1, late fusion and missing blocks; S2, external evaluation. Notes: 1, studies of cross-cancer survival learning and their released code; 2, models and evaluation methods; 3, development and analysis chronology. Citation numbers match the main manuscript.

### Supplementary Table S1 Cohorts, block availability and fold sizes

(a) TCGA target cancers. The table reports eligible patients with a disease-specific survival label, disease-specific deaths, block availability and clinical-model variables for the 13 cancers. Test-fold sizes cover the 25 random folds (five repeats of five folds), counted from the PoE-VAE predictions under natural availability. Blocks with fewer than 50 samples were excluded during preparation: microRNA in GBM and methylation in OV. Every patient in these cancers therefore counts as lacking the excluded block, even if a measurement exists in the source data.

Clinical-variable eligibility. Age was always included. Sex was included when both recorded categories were present and the less common category represented at least 5% of patients with recorded sex. Pathologic stage was used when valid for at least half of eligible patients; otherwise, clinical stage was used if it met that threshold. Grade was included when at least half of patients had a valid value and at least two grades occurred.

(b) Site-held-out folds. Ranges cover five seeds of five folds. The number of distinct partitions was determined by comparing the patients assigned to each fold. When sites are few and large, the assignment rule can produce the same partition for several seeds.

(c) External cohorts. The six CPTAC-3 cohorts were evaluated for overall survival. Eligible patients had at least one block from a primary tumour sample, recorded vital status and a positive follow-up time. Available model inputs were restricted to the blocks used for the corresponding TCGA cancer, including RNA and methylation for GBM. Methylation-feature counts show how many of the 4,000 TCGA-selected features were retained by the cohort-adapted pipeline. Counts in parentheses show how many of those TCGA-selected features were present on the external array. The fully frozen pipeline retained the full TCGA feature set and imputed unavailable features within observed blocks using TCGA training medians.

Recorded sex agreed with RNA and methylation sex markers in every cohort. Areas under the curve were 0.99 to 1.00 for RNA and 0.90 to 1.00 for methylation, against a threshold of 0.90 specified before the check. No patient was excluded on this basis.

| (a) Cancer | Patients | Deaths | All three blocks | Lacking RNA | Lacking microRNA | Lacking methylation | Test patients per fold | Test deaths per fold | Clinical model |
|---|---|---|---|---|---|---|---|---|---|
| BLCA | 396 | 123 | 388 (98.0%) | 6 (1.5%) | 2 (0.5%) | 0 (0.0%) | 79 to 80 | 24 to 25 | Age, sex, pathologic stage, grade |
| BRCA | 1063 | 83 | 758 (71.3%) | 2 (0.2%) | 19 (1.8%) | 293 (27.6%) | 212 to 213 | 16 to 17 | Age, pathologic stage |
| GBM | 223 | 159 | 0 (0.0%) | 81 (36.3%) | 223 (100.0%) | 94 (42.2%) | 44 to 45 | 31 to 32 | Age, sex |
| HNSC | 501 | 130 | 488 (97.4%) | 8 (1.6%) | 5 (1.0%) | 0 (0.0%) | 99 to 101 | 26 to 26 | Age, sex, pathologic stage, grade |
| KIRC | 521 | 110 | 307 (58.9%) | 1 (0.2%) | 19 (3.6%) | 213 (40.9%) | 103 to 105 | 21 to 23 | Age, sex, pathologic stage, grade |
| LGG | 503 | 113 | 499 (99.2%) | 0 (0.0%) | 4 (0.8%) | 0 (0.0%) | 99 to 102 | 22 to 23 | Age, sex, grade |
| LIHC | 362 | 80 | 353 (97.5%) | 5 (1.4%) | 4 (1.1%) | 0 (0.0%) | 72 to 73 | 16 to 16 | Age, sex, pathologic stage, grade |
| LUAD | 471 | 113 | 414 (87.9%) | 3 (0.6%) | 6 (1.3%) | 52 (11.0%) | 93 to 95 | 22 to 23 | Age, sex, pathologic stage |
| LUSC | 445 | 89 | 316 (71.0%) | 3 (0.7%) | 24 (5.4%) | 125 (28.1%) | 89 to 89 | 17 to 18 | Age, sex, pathologic stage |
| OV | 455 | 267 | 0 (0.0%) | 65 (14.3%) | 3 (0.7%) | 455 (100.0%) | 90 to 92 | 53 to 54 | Age, clinical stage, grade |
| PAAD | 176 | 78 | 171 (97.2%) | 5 (2.8%) | 5 (2.8%) | 0 (0.0%) | 35 to 36 | 14 to 16 | Age, sex, pathologic stage, grade |

| (a) Cancer | Patients | Deaths | All three blocks | Lacking RNA | Lacking microRNA | Lacking methylation | Test patients per fold | Test deaths per fold | Clinical model |
|---|---|---|---|---|---|---|---|---|---|
| SARC | 255 | 81 | 251 (98.4%) | 2 (0.8%) | 2 (0.8%) | 0 (0.0%) | 50 to 52 | 16 to 17 | Age, sex |
| STAD | 390 | 100 | 352 (90.3%) | 25 (6.4%) | 7 (1.8%) | 13 (3.3%) | 77 to 79 | 19 to 21 | Age, sex, pathologic stage, grade |
| All | 5761 | 1526 | 4297 (74.6%) | 206 (3.6%) | 323 (5.6%) | 1245 (21.6%) | | | |

| (b) Cancer | Sites | Largest site share | Test patients per fold | Test events per fold | Distinct partitions among the five seeds |
|---|---|---|---|---|---|
| BLCA | 35 | 12.9% | 53 to 111 | 24 to 25 | 5 |
| BRCA | 40 | 13.2% | 56 to 310 | 15 to 21 | 5 |
| GBM | 20 | 35.9% | 28 to 80 | 25 to 59 | 5 |
| HNSC | 28 | 24.2% | 55 to 130 | 20 to 48 | 5 |
| KIRC | 20 | 27.1% | 39 to 168 | 10 to 45 | 1 |
| LGG | 26 | 20.7% | 27 to 233 | 15 to 39 | 4 |
| LIHC | 37 | 40.3% | 24 to 146 | 12 to 28 | 5 |
| LUAD | 33 | 19.3% | 43 to 143 | 22 to 23 | 5 |
| LUSC | 33 | 11.2% | 64 to 142 | 17 to 18 | 5 |
| OV | 20 | 21.1% | 74 to 120 | 48 to 63 | 3 |
| PAAD | 27 | 21.0% | 18 to 56 | 14 to 19 | 5 |
| SARC | 34 | 45.5% | 17 to 116 | 11 to 36 | 5 |
| STAD | 21 | 27.7% | 64 to 108 | 17 to 32 | 2 |

| (c) Cohort | Cases meeting the diagnosis filter | Eligible patients | Deaths | Deaths recorded as cancer-related | Median follow-up of living patients (days) | All blocks used | Lacking RNA | Lacking microRNA | Lacking methylation | Methylation features |
|---|---|---|---|---|---|---|---|---|---|---|
| KIRC | 223 | 199 | 32 | 19 | 1080 | 198 | 0 | 1 | 0 | 3,587 (3,655) |
| LUAD | 230 | 205 | 54 | 25 | 444 | 204 | 0 | 0 | 1 | 3,540 (3,661) |
| LUSC | 109 | 103 | 32 | 15 | 1732 | 101 | 0 | 1 | 1 | 3,557 (3,653) |
| GBM | 202 | 192 | 145 | 106 | 744 | 183 | 4 | not used | 5 | 3,557 (3,644) |
| PAAD | 163 | 160 | 129 | 89 | 550 | 138 | 2 | 17 | 20 | 3,421 (3,658) |
| HNSC | 110 | 106 | 39 | 15 | 907 | 105 | 0 | 1 | 0 | 3,623 (3,658) |

| (c) Cohort | Cases meeting the diagnosis filter | Eligible patients | Deaths | Deaths recorded as cancer-related | Median follow-up of living patients (days) | All blocks used | Lacking RNA | Lacking microRNA | Lacking methylation | Methylation features |
|---|---|---|---|---|---|---|---|---|---|---|
| All | 1037 | 965 | 431 | 269 | | | | | | |

#### Supplementary Table S2 Model selection and tuning budget of each model

Hyperparameters selected within each outer-training set, and earlier exposure of each configuration to the data analysed here. CV, cross-validation.

| Model | Tuned in each training set | Earlier exposure to these data |
|---|---|---|
| PoE-VAE, fixed configuration | None; fixed from earlier work | Architecture and optimiser settings chosen on KIRC and HNSC (overall survival) |
| PoE-VAE, tuned | 20 candidates by inner five-fold CV: 10, 25, 50, 100 or 200 training steps; encoder width and latent dimension (64, 16) or (128, 48); Kullback–Leibler weight 0.001 or 0.1 | Candidate grid specified after the earlier analyses were reported; it contains the fixed configuration |
| Borrowing model | Probe ridge penalty (11 values) and fusion weight (11 values), inner five-fold CV | Recipe chosen after several transfer recipes were tried on KIRC and HNSC |
| Elastic-net late fusion | Penalty per block, path of 30 values, inner five-fold CV; mixing parameter fixed at 0.5 | None |
| Elastic-net concatenation | Penalty, path of 30 values, inner five-fold CV; mixing parameter fixed at 0.5 | None |
| Neural late fusion, fixed configuration | None; 100 steps with the optimiser settings of the PoE-VAE | None |
| Neural late fusion, tuned | 12 candidates per block by inner five-fold CV: 10, 25, 50, 100, 200 or 400 steps; hidden width 32 or 128 | Candidate grid specified after the earlier analyses were reported; it contains the fixed configuration |
| PriorityLasso | Penalty per block by inner five-fold CV; block order by cross-validated single-block concordance | One configuration (offset imputation, zero for missing test blocks) |
| BlockForest | Block weights: 50 candidate sets of 500 trees (package default 300 sets of 1,500); final forest 1,000 trees | One configuration (mean-filled blocks) |
| Clinical model | None; ridge penalty fixed at 0.1 | Variables by a fixed availability rule |
| VAECox reimplementation, pretrained and randomly initialised networks (both data settings) | Six candidates by inner five-fold CV: learning rate $10^{-4}$ or $10^{-3}$ by weight decay $10^{-3}$, $10^{-4}$ or $10^{-5}$, each scored by the maximum over the 21 fine-tuning steps of its mean validation concordance. Training length: fixed at 21 steps, or, in a follow-up, the step at which the chosen candidate's mean validation concordance is highest | None; architecture, search grid and numbers of steps taken from the published code |

The fixed configuration of the PoE-VAE was chosen in 3.4% of partitions. Chosen training steps for the PoE-VAE: 10 in 40%, 25 in 20%, 50 in 11%, 100 in 11% and 200 in 18% of partitions; for the block models of neural late fusion: 10 in 46%, 25 in 18%, 50 in 9%, 100 in 5%, 200 in 7% and 400 in 15%. Where the fixed configurations were refitted, test predictions equalled those of the fixed PoE-VAE and neural late fusion (largest absolute difference 0).

#### Supplementary Table S3 Concordance, borrowing gains and integrated Brier scores per cancer

Disease-specific survival in 13 TCGA cancers; five repeats of five-fold cross-validation; means under natural block availability. (a) Harrell's concordance; the two tuned models choose their hyperparameters by inner cross-validation, and the other models use the configurations in Supplementary Table S2. (b) Uno's concordance. (c) Borrowing model minus the PoE-VAE (Harrell's concordance), with 95% paired patient-bootstrap intervals, random-effects pooling weights, fusion weights selected on the training data and the bootstrap standard deviation (SD); cancers are ordered by pooling weight. (d) Integrated Brier scores, for which lower is better. PriorityLasso has no Brier score. Risk-score models use cross-fitted mappings, BlockForest its own curves, and Kaplan–Meier is the training-only reference.

| (a) Cancer | PoE-VAE | Borrowing model | Elastic-net late fusion | Elastic-net concatenation | Neural late fusion | PriorityLasso | BlockForest | Clinical model | Tuned PoE-VAE | Tuned neural late fusion |
|---|---|---|---|---|---|---|---|---|---|---|
| BLCA | 0.658 | 0.671 | 0.638 | 0.604 | 0.681 | 0.611 | 0.651 | 0.679 | 0.660 | 0.679 |
| BRCA | 0.697 | 0.712 | 0.682 | 0.666 | 0.716 | 0.607 | 0.647 | 0.757 | 0.714 | 0.708 |
| GBM | 0.565 | 0.561 | 0.550 | 0.528 | 0.553 | 0.546 | 0.528 | 0.630 | 0.559 | 0.557 |
| HNSC | 0.622 | 0.647 | 0.635 | 0.610 | 0.643 | 0.651 | 0.627 | 0.582 | 0.622 | 0.643 |
| KIRC | 0.786 | 0.787 | 0.765 | 0.746 | 0.772 | 0.741 | 0.757 | 0.841 | 0.779 | 0.775 |
| LGG | 0.836 | 0.840 | 0.848 | 0.850 | 0.837 | 0.847 | 0.826 | 0.783 | 0.832 | 0.843 |
| LIHC | 0.685 | 0.701 | 0.712 | 0.677 | 0.716 | 0.705 | 0.709 | 0.698 | 0.704 | 0.720 |
| LUAD | 0.606 | 0.639 | 0.616 | 0.586 | 0.610 | 0.601 | 0.626 | 0.662 | 0.604 | 0.620 |
| LUSC | 0.577 | 0.583 | 0.574 | 0.565 | 0.585 | 0.568 | 0.602 | 0.641 | 0.603 | 0.593 |
| OV | 0.554 | 0.549 | 0.548 | 0.535 | 0.568 | 0.543 | 0.550 | 0.606 | 0.583 | 0.578 |
| PAAD | 0.601 | 0.609 | 0.591 | 0.599 | 0.582 | 0.625 | 0.602 | 0.591 | 0.596 | 0.615 |
| SARC | 0.657 | 0.650 | 0.665 | 0.662 | 0.670 | 0.645 | 0.650 | 0.564 | 0.661 | 0.663 |
| STAD | 0.591 | 0.594 | 0.585 | 0.568 | 0.588 | 0.571 | 0.562 | 0.638 | 0.578 | 0.577 |

| (b) Cancer | PoE-VAE | Borrowing model | Elastic-net late fusion | Elastic-net concatenation | Neural late fusion | PriorityLasso | BlockForest | Clinical model |
|---|---|---|---|---|---|---|---|---|
| BLCA | 0.652 | 0.659 | 0.625 | 0.596 | 0.677 | 0.602 | 0.642 | 0.675 |
| BRCA | 0.645 | 0.664 | 0.632 | 0.628 | 0.662 | 0.578 | 0.576 | 0.684 |
| GBM | 0.558 | 0.555 | 0.546 | 0.525 | 0.549 | 0.542 | 0.528 | 0.634 |
| HNSC | 0.622 | 0.645 | 0.642 | 0.619 | 0.641 | 0.656 | 0.630 | 0.567 |
| KIRC | 0.776 | 0.777 | 0.759 | 0.736 | 0.765 | 0.737 | 0.745 | 0.832 |
| LGG | 0.789 | 0.786 | 0.797 | 0.798 | 0.782 | 0.788 | 0.766 | 0.740 |
| LIHC | 0.688 | 0.682 | 0.672 | 0.636 | 0.699 | 0.655 | 0.680 | 0.725 |
| LUAD | 0.579 | 0.633 | 0.606 | 0.573 | 0.597 | 0.581 | 0.582 | 0.654 |
| LUSC | 0.558 | 0.569 | 0.556 | 0.549 | 0.562 | 0.553 | 0.584 | 0.662 |
| OV | 0.556 | 0.551 | 0.549 | 0.534 | 0.566 | 0.542 | 0.544 | 0.596 |
| PAAD | 0.602 | 0.612 | 0.591 | 0.591 | 0.587 | 0.626 | 0.605 | 0.594 |
| SARC | 0.651 | 0.645 | 0.664 | 0.665 | 0.672 | 0.646 | 0.648 | 0.561 |
| STAD | 0.593 | 0.592 | 0.584 | 0.567 | 0.583 | 0.571 | 0.559 | 0.626 |

| (c) Cancer | Gain | 95% interval | Weight | Mean fusion weight | Bootstrap SD |
|---|---|---|---|---|---|
| KIRC | +0.0013 | (−0.0069, +0.0101) | 26.9% | 0.24 | 0.0045 |

| (c) Cancer | Gain | 95% interval | Weight | Mean fusion weight | Bootstrap SD |
|---|---|---|---|---|---|
| LGG | +0.0041 | (−0.0050, +0.0142) | 22.5% | 0.28 | 0.0049 |
| GBM | −0.0039 | (−0.0143, +0.0068) | 17.8% | 0.16 | 0.0055 |
| OV | −0.0044 | (−0.0175, +0.0079) | 12.8% | 0.24 | 0.0065 |
| SARC | −0.0070 | (−0.0322, +0.0200) | 3.1% | 0.39 | 0.0133 |
| STAD | +0.0030 | (−0.0229, +0.0307) | 3.0% | 0.35 | 0.0135 |
| BLCA | +0.0132 | (−0.0134, +0.0398) | 2.9% | 0.39 | 0.0137 |
| LIHC | +0.0158 | (−0.0117, +0.0417) | 2.8% | 0.37 | 0.0140 |
| LUSC | +0.0064 | (−0.0240, +0.0361) | 2.3% | 0.49 | 0.0153 |
| BRCA | +0.0146 | (−0.0159, +0.0445) | 2.2% | 0.31 | 0.0157 |
| HNSC | +0.0255 | (−0.0063, +0.0584) | 1.9% | 0.60 | 0.0169 |
| LUAD | +0.0333 | (−0.0127, +0.0803) | 1.0% | 0.73 | 0.0238 |
| PAAD | +0.0087 | (−0.0388, +0.0574) | 0.9% | 0.53 | 0.0243 |

| (d) Cancer | PoE-VAE | Borrowing model | Elastic-net late fusion | Elastic-net concatenation | Neural late fusion | BlockForest | Clinical model | Kaplan–Meier |
|---|---|---|---|---|---|---|---|---|
| BLCA | 0.1300 | 0.1286 | 0.1315 | 0.1353 | 0.1248 | 0.1330 | 0.1296 | 0.1384 |
| BRCA | 0.0641 | 0.0645 | 0.0646 | 0.0649 | 0.0634 | 0.0664 | 0.0605 | 0.0666 |
| GBM | 0.2079 | 0.2083 | 0.2079 | 0.2102 | 0.2071 | 0.2099 | 0.1924 | 0.2095 |
| HNSC | 0.1172 | 0.1146 | 0.1166 | 0.1183 | 0.1144 | 0.1180 | 0.1210 | 0.1216 |
| KIRC | 0.0910 | 0.0915 | 0.0958 | 0.0999 | 0.0942 | 0.0977 | 0.0841 | 0.1098 |
| LGG | 0.1075 | 0.1056 | 0.1060 | 0.1044 | 0.1047 | 0.1102 | 0.1239 | 0.1547 |
| LIHC | 0.1347 | 0.1352 | 0.1331 | 0.1382 | 0.1307 | 0.1331 | 0.1328 | 0.1458 |
| LUAD | 0.1239 | 0.1210 | 0.1226 | 0.1246 | 0.1214 | 0.1236 | 0.1216 | 0.1260 |
| LUSC | 0.1148 | 0.1143 | 0.1157 | 0.1151 | 0.1137 | 0.1136 | 0.1128 | 0.1158 |
| OV | 0.1906 | 0.1909 | 0.1902 | 0.1918 | 0.1880 | 0.1903 | 0.1873 | 0.1918 |
| PAAD | 0.1613 | 0.1600 | 0.1594 | 0.1593 | 0.1607 | 0.1602 | 0.1611 | 0.1634 |
| SARC | 0.1409 | 0.1422 | 0.1386 | 0.1421 | 0.1371 | 0.1441 | 0.1524 | 0.1526 |
| STAD | 0.1220 | 0.1216 | 0.1235 | 0.1232 | 0.1222 | 0.1254 | 0.1212 | 0.1236 |

#### Supplementary Table S4 Pooled paired contrasts and sensitivity analyses

(a) Concordance contrasts. Values are changes in Harrell's concordance, first-named minus second-named, pooled over the 13 TCGA cancers under natural block availability unless stated otherwise. Masked analyses use complete-case test patients from 11 cancers under the seven block-availability patterns. A dash indicates that a quantity was not computed. The variants in the last group are defined in Supplementary Note 2. Intervals for test-selected quantities condition on the selection.

Pooling and intervals. Random-effects summaries use Hartung–Knapp intervals at the level shown, assigned within each analysis family (Supplementary Note 3). The modified Hartung–Knapp method [46] changes the interval but leaves the estimate unchanged. The arithmetic mean gives equal weight to each cancer, and its percentile interval treats these cancers as fixed. Each contrast has its own pooling weights, so separately pooled contrasts need not add up.

Endpoint sensitivity. The ten-cancer summary excludes BRCA, LGG and LIHC, for which the curators caution against disease-specific survival [6]. It re-pools the same per-cancer estimates using the same random-effects estimator and interval levels; no models were refitted. The modified intervals, arithmetic means and ten-cancer summaries are post hoc.

Zero bootstrap variance. A cancer with zero bootstrap variance was assigned the smallest positive variance observed for that contrast. This was required only for the best of three borrowing models against the borrowing model, where the models coincided in BLCA, GBM, KIRC and LGG. Excluding these four cancers gave a random-effects summary of +0.0059 (+0.0035, +0.0084).

(b) Integrated Brier scores. Comparisons with the Kaplan–Meier reference report model minus Kaplan–Meier, pooled by random effects with nominal 95% intervals. Relative improvement, (Kaplan–Meier − model) / Kaplan–Meier, is averaged over the 13 cancers with equal weight. For contrasts between models, a negative value favours the first-named model. Uno's concordance comparisons are reported in Table 3 of the main manuscript.

| **(a) Contrast** | **Level** | **Random-effects summary** | **Modified Hartung–Knapp interval** | **Arithmetic mean across cancers** | **Ten cancers without an endpoint caution, random-effects summary** |
|---|---|---|---|---|---|
| *Borrowing* | | | | | |
| Borrowing model − PoE-VAE | 95% | +0.0021 (−0.0023, +0.0064) | (−0.0030, +0.0072) | +0.0085 (+0.0006, +0.0164) | +0.0005 (−0.0048, +0.0059) |
| *Model comparisons, fixed configurations* | | | | | |
| PoE-VAE − elastic-net late fusion | 97.5% | +0.0015 (−0.0093, +0.0122) | (−0.0096, +0.0125) | +0.0019 (−0.0095, +0.0131) | +0.0067 (−0.0043, +0.0178) |
| PoE-VAE − elastic-net concatenation | 97.5% | +0.0180 (+0.0034, +0.0327) | (+0.0011, +0.0350) | +0.0183 (+0.0055, +0.0308) | +0.0258 (+0.0109, +0.0406) |
| Elastic-net late fusion − elastic-net concatenation | 95% | +0.0160 (+0.0082, +0.0239) | (+0.0077, +0.0244) | +0.0164 (+0.0096, +0.0235) | – |
| PoE-VAE − PriorityLasso | 97.5% | +0.0143 (−0.0092, +0.0379) | (−0.0098, +0.0385) | +0.0133 (+0.0005, +0.0261) | +0.0141 (−0.0077, +0.0359) |

| (a) Contrast | Level | Random-effects summary | Modified Hartung–Knapp interval | Arithmetic mean across cancers | Ten cancers without an endpoint caution, random-effects summary |
|---|---|---|---|---|---|
| PoE-VAE − neural late fusion | 97.5% | −0.0053 (−0.0159, +0.0053) | (−0.0161, +0.0055) | −0.0066 (−0.0167, +0.0040) | −0.0040 (−0.0171, +0.0091) |
| Neural late fusion − elastic-net late fusion | 95% | +0.0079 (−0.0021, +0.0178) | (−0.0026, +0.0183) | +0.0085 (−0.0011, +0.0182) | – |
| PoE-VAE − BlockForest | 97.5% | +0.0080 (−0.0057, +0.0218) | – | – | – |
| Elastic-net late fusion − BlockForest | 97.5% | +0.0082 (−0.0024, +0.0189) | – | – | – |
| PoE-VAE − clinical model | 97.5% | −0.0196 (−0.0555, +0.0162) | – | – | – |
| Elastic-net late fusion − clinical model | 97.5% | −0.0202 (−0.0630, +0.0225) | – | – | – |
| *Models with hyperparameters chosen by inner cross-validation* | | | | | |
| Tuned PoE-VAE − elastic-net late fusion | 97.5% | +0.0058 (−0.0076, +0.0192) | (−0.0085, +0.0201) | +0.0064 (−0.0034, +0.0167) | – |
| Tuned neural late fusion − elastic-net late fusion | 97.5% | +0.0116 (+0.0010, +0.0222) | (+0.0002, +0.0230) | +0.0125 (+0.0027, +0.0220) | – |
| Tuned PoE-VAE − fixed PoE-VAE | 95% | +0.0015 (−0.0059, +0.0089) | – | +0.0046 (−0.0007, +0.0101) | – |
| Tuned neural late fusion − fixed neural late fusion | 95% | +0.0023 (−0.0021, +0.0066) | – | +0.0040 (−0.0009, +0.0092) | – |
| Tuned PoE-VAE − tuned neural late fusion | 95% | −0.0065 (−0.0130, −0.0001) | – | −0.0060 (−0.0128, +0.0013) | – |
| Tuned PoE-VAE − PriorityLasso | 95% | +0.0182 (−0.0040, +0.0403) | – | +0.0178 (+0.0079, +0.0282) | – |
| Tuned PoE-VAE − elastic-net late fusion, masked | 95% | +0.0050 (−0.0068, +0.0167) | – | +0.0069 (−0.0009, +0.0154) | – |
| Tuned neural late fusion − elastic-net late fusion, masked | 95% | +0.0128 (+0.0032, +0.0224) | – | +0.0141 (+0.0057, +0.0221) | – |
| *Features selected in the target cancer instead of the other cohorts* | | | | | |
| PoE-VAE, target-selected − shared features | 97.5% | −0.0001 (−0.0137, +0.0135) | – | – | +0.0048 (−0.0078, +0.0174) |

| (a) Contrast | Level | Random-effects summary | Modified Hartung–Knapp interval | Arithmetic mean across cancers | Ten cancers without an endpoint caution, random-effects summary |
|---|---|---|---|---|---|
| Elastic-net late fusion, target-selected − shared features | 97.5% | +0.0024 (−0.0072, +0.0121) | – | – | +0.0015 (−0.0088, +0.0117) |
| *Site-held-out folds* | | | | | |
| PoE-VAE, site-held-out − random folds | 97.5% | −0.0229 (−0.0400, −0.0058) | (−0.0400, −0.0058) | −0.0259 (−0.0392, −0.0138) | – |
| Elastic-net late fusion, site-held-out − random folds | 97.5% | −0.0156 (−0.0291, −0.0021) | (−0.0291, −0.0021) | −0.0183 (−0.0310, −0.0055) | – |
| PoE-VAE − elastic-net late fusion, site-held-out folds | 95% | −0.0043 (−0.0177, +0.0092) | (−0.0180, +0.0095) | −0.0058 (−0.0182, +0.0064) | – |
| *Departures from the reference evaluation of borrowing* | | | | | |
| Borrowing with test-selected weight − PoE-VAE | 95% | +0.0066 (+0.0027, +0.0105) | (+0.0026, +0.0106) | +0.0169 (+0.0090, +0.0245) | – |
| Borrowing with test-selected weight − borrowing model | 95% | +0.0057 (+0.0030, +0.0084) | (+0.0028, +0.0086) | +0.0084 (+0.0051, +0.0113) | – |
| Best of three borrowing models (test-selected) − PoE-VAE | 95% | +0.0034 (−0.0018, +0.0086) | (−0.0021, +0.0089) | +0.0125 (+0.0029, +0.0219) | – |
| Best of three borrowing models (test-selected) − borrowing model | 95% | +0.0024 (+0.0002, +0.0046) | (−0.0004, +0.0053) | +0.0040 (+0.0007, +0.0073) | – |
| Borrowing with target-inclusive pretraining − PoE-VAE | 95% | +0.0052 (+0.0010, +0.0094) | (−0.0002, +0.0106) | +0.0085 (+0.0004, +0.0161) | – |
| Borrowing with target-inclusive pretraining − borrowing model | 95% | +0.0009 (−0.0014, +0.0032) | (−0.0016, +0.0034) | −0.0000 (−0.0033, +0.0031) | – |
| Joint training across cancers − PoE-VAE | 95% | −0.0115 (−0.0303, +0.0073) | (−0.0303, +0.0073) | −0.0113 (−0.0251, +0.0031) | – |

| (b) Integrated Brier score | Pooled difference (95% interval) | Mean relative improvement |
|---|---|---|
| *Model minus the Kaplan–Meier reference* | | |
| PoE-VAE | −0.0054 (−0.0104, −0.0004) | 6.4% |
| Borrowing model | −0.0061 (−0.0113, −0.0010) | 6.8% |
| Elastic-net late fusion | −0.0050 (−0.0097, −0.0002) | 6.2% |

| (b) Integrated Brier score | Pooled difference (95% interval) | Mean relative improvement |
|---|---|---|
| Elastic-net concatenation | −0.0033 (−0.0076, +0.0010) | 5.0% |
| Neural late fusion | −0.0070 (−0.0123, −0.0016) | 7.6% |
| BlockForest | −0.0054 (−0.0112, +0.0004) | 5.1% |
| Clinical model | −0.0060 (−0.0104, −0.0016) | 6.8% |
| *Contrasts between models* | | |
| PoE-VAE − elastic-net concatenation (post hoc) | −0.0013 (−0.0026, −0.0001) | – |
| PoE-VAE − elastic-net late fusion | −0.0003 (−0.0010, +0.0004) | – |
| PoE-VAE − neural late fusion | +0.0014 (+0.0002, +0.0025) | – |
| PoE-VAE − BlockForest | −0.0015 (−0.0028, −0.0003) | – |
| PoE-VAE − clinical model | +0.0009 (−0.0027, +0.0045) | – |
| Borrowing model − PoE-VAE | −0.0001 (−0.0006, +0.0004) | – |
| Elastic-net late fusion − elastic-net concatenation | −0.0015 (−0.0026, −0.0003) | – |

#### Supplementary Table S5 VAECox reimplementation: four versions of the evaluation procedure, unequal test-step selection and validation-selected steps

The table reports differences in Harrell's concordance between the pretrained and randomly initialised networks in our VAECox reimplementation [10], evaluated on test patients with RNA data. Differences are pretrained minus randomly initialised, with 95% intervals. Rows pooled by random effects have their own weights, so separately pooled rows need not add up.

The published-style procedure pretrains on an 80% sample of the pooled patients from all 20 cohorts, including target-cancer test patients. It scales using all target patients and evaluates each network at its best test step. The original study reported a gain of +0.020. Supplementary Note 2 lists differences from that study.

(a) Thirteen-cancer analysis: disease-specific survival, 4,000 genes per target and a random-effects summary.

(b) Ten-cancer overall-survival analysis: VAECox's ten target cancers and 20 pretraining cancers, with 18,858 genes. This setting is exploratory.

(c) Unequal test-step selection: the same fitted networks and test patients are used to compare selection of the best test step for either network, both networks or neither. The 95% intervals condition on the selected steps. This post hoc analysis was specified after its point estimates had been inspected. Because the best test step cannot score below the last step on the same test set, the informative quantity is the size of the shift. The published code evaluates every network at its best test step; the two unequal-selection rows are a constructed demonstration and do not describe the original study.

(d) Validation-selected evaluation: the target-excluded protocol with each network evaluated at its validation-selected step. This exploratory follow-up is described below the panel.

| **(a) Version** | **13 cancers** | **The ten of these 13 that VAECox studied** |
|---|---|---|
| Published-style procedure: both networks evaluated at the best test step | −0.0043 (−0.0099, +0.0014) | −0.0051 (−0.0118, +0.0017) |
| Both networks evaluated at the last step | −0.0054 (−0.0123, +0.0016) | −0.0076 (−0.0156, +0.0005) |
| No test-patient data in pretraining or scaling | −0.0052 (−0.0123, +0.0018) | −0.0068 (−0.0159, +0.0023) |
| Target-excluded protocol: target cancer excluded from pretraining | −0.0043 (−0.0113, +0.0027) | −0.0051 (−0.0139, +0.0037) |
| Effect of test-step selection (row 1 − row 2) | +0.0033 (−0.0008, +0.0074) | +0.0039 (−0.0004, +0.0083) |
| Effect of test patients' data (row 2 − row 3) | −0.0004 (−0.0050, +0.0041) | −0.0006 (−0.0060, +0.0048) |
| Effect of the target cancer in pretraining (row 3 − row 4) | −0.0014 (−0.0059, +0.0032) | −0.0019 (−0.0069, +0.0031) |

The published-style summary for (a) weights VAECox's ten cancers by patient count and has no interval. At the best test step, concordance was 0.7126 for the pretrained network and 0.7181 for the randomly initialised network. At the last step, the corresponding values were 0.6652 and 0.6754.

With equal weight across the 13 cancers, choosing the best test step raised mean concordance by 0.049 for the pretrained network and 0.047 for the randomly initialised network. In individual cancers, the increases ranged from 0.009 to 0.092. Under the target-excluded protocol, KIRC was the only cancer whose interval excluded zero (difference −0.025, interval −0.037 to −0.014).

| **(b) Version** | **Random-effects summary** | **Arithmetic mean across cancers** |
|---|---|---|
| Published-style procedure: both networks evaluated at the best test step | −0.0057 (−0.0107, −0.0007) | −0.0048 (−0.0104, +0.0010) |
| Both networks evaluated at the last step | −0.0059 (−0.0134, +0.0016) | −0.0054 (−0.0108, −0.0001) |
| No test-patient data in pretraining or scaling | −0.0080 (−0.0139, −0.0022) | −0.0080 (−0.0135, −0.0026) |
| Target-excluded protocol: target cancer excluded from pretraining | −0.0092 (−0.0158, −0.0025) | −0.0092 (−0.0150, −0.0035) |
| Effect of test-step selection (row 1 − row 2) | +0.0008 (−0.0036, +0.0051) | +0.0005 (−0.0046, +0.0062) |
| Effect of test patients' data (row 2 − row 3) | +0.0026 (−0.0028, +0.0080) | +0.0026 (−0.0005, +0.0055) |
| Effect of the target cancer in pretraining (row 3 − row 4) | +0.0009 (−0.0053, +0.0071) | +0.0012 (−0.0018, +0.0040) |

The published-style summary for (b), weighted by patient count across the ten cancers, gave concordance of 0.6713 for the pretrained network and 0.6776 for the randomly initialised network at the best test step. At the last step, the corresponding values were 0.6443 and 0.6516. The modified Hartung–Knapp interval for the published-style procedure was −0.0119 to +0.0006.



With equal weight across cancers, choosing the best test step raised mean concordance by 0.027 for both networks; increases within individual cancers ranged from 0.008 to 0.059. Under the target-excluded protocol, BRCA was the only cancer whose interval excluded zero (difference −0.033, interval −0.055 to −0.008). In LUSC, overall-survival concordance was below 0.5 for every version (0.47 to 0.48).

| **(c) Setting** | **Evaluated at the best test step** | **Random-effects summary** | **Arithmetic mean across cancers** | **Cancers with a positive difference** |
|---|---|---|---|---|
| 13 cancers, disease-specific survival | Neither network | −0.0054 (−0.0123, +0.0016) | −0.0040 (−0.0094, +0.0011) | 5 of 13 |
| 13 cancers, disease-specific survival | Both networks | −0.0043 (−0.0099, +0.0014) | −0.0019 (−0.0069, +0.0032) | 6 of 13 |
| 13 cancers, disease-specific survival | Pretrained network only | +0.0405 (+0.0220, +0.0590) | +0.0448 (+0.0353, +0.0540) | 12 of 13 |
| 13 cancers, disease-specific survival | Randomly initialised network only | −0.0444 (−0.0556, −0.0331) | −0.0508 (−0.0593, −0.0423) | 0 of 13 |
| Ten cancers, overall survival | Neither network | −0.0059 (−0.0134, +0.0016) | −0.0054 (−0.0108, −0.0001) | 2 of 10 |
| Ten cancers, overall survival | Both networks | −0.0057 (−0.0107, −0.0007) | −0.0048 (−0.0104, +0.0010) | 1 of 10 |
| Ten cancers, overall survival | Pretrained network only | +0.0176 (+0.0064, +0.0288) | +0.0220 (+0.0141, +0.0302) | 9 of 10 |
| Ten cancers, overall survival | Randomly initialised network only | −0.0289 (−0.0401, −0.0177) | −0.0322 (−0.0394, −0.0246) | 0 of 10 |

For the row selecting the best test step only for the pretrained network, modified Hartung–Knapp intervals were +0.0214 to +0.0596 in the 13-cancer analysis and +0.0061 to +0.0291 in the ten-cancer setting.

Relative to evaluating both networks at the last step, selecting the best test step only for the pretrained network shifted the arithmetic mean difference by +0.0489 in the 13 cancers and +0.0273 in the ten-cancer setting. Selecting only for the randomly initialised network shifted it by −0.0467 and −0.0268, respectively. These shifts are point estimates.

All four rows within each setting of panel (c) use the same fitted networks. The two unequal-selection rows are therefore not independent evidence. They measure a shift from a fixed last-step rule, not the full optimism relative to independent data.

| **(d) Setting** | **Pretrained minus randomly initialised network** | **Random-effects summary** | **Modified Hartung–Knapp interval** | **Arithmetic mean across cancers** |
|---|---|---|---|---|
| 13 cancers, disease-specific survival | Last step (repeat of panel a or b) | −0.0043 (−0.0113, +0.0027) | (−0.0114, +0.0028) | −0.0033 (−0.0082, +0.0016) |
| 13 cancers, disease-specific survival | Validation-selected step | −0.0007 (−0.0117, +0.0104) | (−0.0118, +0.0104) | +0.0001 (−0.0049, +0.0056) |
| 13 cancers, disease-specific survival | Change, validation-selected minus last step | +0.0027 (−0.0051, +0.0104) | (−0.0051, +0.0104) | +0.0035 (−0.0021, +0.0091) |
| Ten cancers, overall survival | Last step (repeat of panel a or b) | −0.0092 (−0.0158, −0.0025) | (−0.0158, −0.0025) | −0.0092 (−0.0150, −0.0035) |
| Ten cancers, overall survival | Validation-selected step | −0.0041 (−0.0079, −0.0002) | (−0.0103, +0.0022) | −0.0043 (−0.0100, +0.0017) |
| Ten cancers, overall survival | Change, validation-selected minus last step | +0.0040 (+0.0005, +0.0075) | (−0.0001, +0.0082) | +0.0049 (+0.0001, +0.0100) |

(d) Validation-selected follow-up. This exploratory analysis was specified on 12 October 2026, after the results above were known but before the follow-up was run. Each network under the target-excluded protocol was evaluated at the step with the highest mean inner validation concordance for its chosen candidate, taking the first step if several tied. The step was selected separately for each network and partition, without test patients. Refitting with the seeds of panels (a) and (b) reproduced the last-step predictions exactly and selected the same candidates.

In the 13-cancer analysis, the validation-selected step was the last step in 45 of 325 partitions for the pretrained network and 32 for the randomly initialised network. In the ten-cancer setting, the corresponding counts were 29 and 24 of 250 partitions. The networks selected the same step in 62 partitions in the 13-cancer analysis and 36 in the ten-cancer setting.

At the validation-selected step, cancer-specific 95% intervals excluded zero for LIHC (+0.030) and LUSC (+0.030), favouring pretraining, and for KIRC (−0.026), HNSC (−0.020) and OV (−0.014), favouring random initialisation. These values are point estimates from the 13-cancer analysis. No cancer-specific interval excluded zero in the ten-cancer setting.

#### Supplementary Table S6 Variation between single runs

Post hoc; specified after the point estimates had been inspected. The difference for each five-fold run, and for each train-test partition, minus the mean of its cancer's 25 partitions; standard deviation and share at least 0.02 from that mean, pooled over cancers. This variation combines the choice of partition with the randomness of model fitting, and deviations are measured against the five-run mean, not a known true value. The standard deviation is the sample standard deviation (denominator n − 1) of all centred values of a contrast. All eleven contrasts that were analysed are listed. Figure 3 of the main manuscript plots the seven contrasts among the multi-omics models.

| Contrast | SD of one five-fold run | Runs at least 0.02 from the mean | SD of one partition | Partitions at least 0.02 from the mean |
|---|---|---|---|---|
| *Multi-omics models, 13 cancers, disease-specific survival* | | | | |
| Borrowing model − PoE-VAE | 0.0122 | 10.8% of 65 | 0.0386 | 45.5% of 325 |
| PoE-VAE − elastic-net late fusion | 0.0148 | 23.1% of 65 | 0.0490 | 64.9% of 325 |
| PoE-VAE − elastic-net concatenation | 0.0192 | 29.2% of 65 | 0.0553 | 71.1% of 325 |
| Elastic-net late fusion − elastic-net concatenation | 0.0165 | 18.5% of 65 | 0.0430 | 53.5% of 325 |
| PoE-VAE − PriorityLasso | 0.0215 | 41.5% of 65 | 0.0609 | 70.2% of 325 |
| PoE-VAE − neural late fusion | 0.0124 | 12.3% of 65 | 0.0454 | 62.2% of 325 |
| Neural late fusion − elastic-net late fusion | 0.0149 | 18.5% of 65 | 0.0479 | 64.6% of 325 |
| *VAECox reimplementation, pretrained − randomly initialised network, 13 cancers, disease-specific survival* | | | | |
| Target-excluded protocol | 0.0195 | 23.1% of 65 | 0.0381 | 50.5% of 325 |
| Published-style procedure | 0.0094 | 6.2% of 65 | 0.0250 | 36.0% of 325 |
| *VAECox reimplementation, pretrained − randomly initialised network, ten cancers, overall survival* | | | | |
| Target-excluded protocol | 0.0092 | 4.0% of 50 | 0.0278 | 44.0% of 250 |
| Published-style procedure | 0.0090 | 0.0% of 50 | 0.0227 | 36.4% of 250 |

#### Supplementary Table S7 Site-held-out and external evaluation

(a) Change in Harrell's concordance per cancer, site-held-out minus random folds, with nominal 95% paired bootstrap intervals; the last column is the change for the PoE-VAE minus the change for elastic-net late fusion. Fold sizes are in Supplementary Table S1 and pooled changes in Supplementary Table S4. (b) Harrell's concordance for overall survival in the six CPTAC-3 cohorts (patients and deaths in Supplementary Table S1), with nominal 95% intervals, and the random-effects summary over the six cohorts. Models were fitted to the corresponding TCGA cancer with disease-specific survival and applied without refitting. Cohort-adapted: features retained by their coverage and variability in the external cohort and scaled with the external cohort's own statistics (the primary external analysis). TCGA scaling: the same features scaled with TCGA training statistics (prespecified sensitivity analysis). Fully frozen: the full TCGA feature set, TCGA scaling statistics and TCGA training medians for features missing from the external data (exploratory; specified after the other two). No pipeline used external outcomes. The three pipelines are applied to the same patients and are not independent validations; the fully frozen models were fitted on a different feature set, so differences between pipelines are descriptive. (c) Paired contrasts pooled over the six cohorts, first-named minus second-named, at the level shown; the TCGA-scaling contrasts have 95% intervals.

| **(a) Cancer** | **PoE-VAE** | **Elastic-net late fusion** | **Difference between the changes** |
|---|---|---|---|
| BLCA | −0.035 (−0.062, −0.011) | −0.015 (−0.042, +0.012) | −0.020 (−0.044, +0.004) |
| BRCA | −0.005 (−0.042, +0.031) | −0.004 (−0.039, +0.033) | −0.001 (−0.039, +0.034) |
| GBM | −0.009 (−0.050, +0.032) | +0.004 (−0.037, +0.049) | −0.013 (−0.061, +0.031) |
| HNSC | −0.043 (−0.077, −0.012) | −0.013 (−0.044, +0.018) | −0.030 (−0.059, −0.004) |
| KIRC | −0.053 (−0.090, −0.020) | −0.061 (−0.102, −0.024) | +0.009 (−0.016, +0.036) |
| LGG | −0.023 (−0.059, +0.007) | −0.024 (−0.050, −0.0002) | +0.001 (−0.026, +0.026) |
| LIHC | −0.107 (−0.177, −0.046) | −0.083 (−0.156, −0.016) | −0.024 (−0.085, +0.039) |
| LUAD | −0.005 (−0.042, +0.029) | −0.015 (−0.047, +0.015) | +0.009 (−0.024, +0.039) |
| LUSC | +0.013 (−0.023, +0.044) | +0.011 (−0.029, +0.049) | +0.002 (−0.032, +0.036) |
| OV | −0.025 (−0.051, +0.002) | −0.018 (−0.045, +0.009) | −0.007 (−0.036, +0.024) |
| PAAD | −0.013 (−0.060, +0.032) | −0.018 (−0.061, +0.029) | +0.005 (−0.047, +0.053) |
| SARC | −0.025 (−0.073, +0.019) | −0.002 (−0.058, +0.045) | −0.023 (−0.069, +0.026) |
| STAD | −0.006 (−0.048, +0.032) | +0.003 (−0.034, +0.040) | −0.008 (−0.046, +0.027) |

| **(b) Model** | **KIRC** | **LUAD** | **LUSC** | **GBM** | **PAAD** | **HNSC** | **Pooled** |
|---|---|---|---|---|---|---|---|
| *Cohort-adapted pipeline* | | | | | | | |
| PoE-VAE | 0.668<br>(0.537, 0.776) | 0.625<br>(0.538, 0.701) | 0.575<br>(0.471, 0.680) | 0.553<br>(0.503, 0.602) | 0.597<br>(0.544, 0.650) | 0.576<br>(0.479, 0.669) | 0.586<br>(0.550, 0.623) |
| Borrowing model | 0.677<br>(0.549, 0.785) | 0.606<br>(0.508, 0.696) | 0.624<br>(0.532, 0.714) | 0.558<br>(0.511, 0.603) | 0.603<br>(0.558, 0.647) | 0.584<br>(0.484, 0.685) | 0.593<br>(0.558, 0.627) |

| (b) Model | KIRC | LUAD | LUSC | GBM | PAAD | HNSC | Pooled |
|---|---|---|---|---|---|---|---|
| Elastic-net late fusion | 0.684 (0.555, 0.791) | 0.639 (0.557, 0.712) | 0.602 (0.503, 0.698) | 0.573 (0.526, 0.619) | 0.584 (0.530, 0.638) | 0.597 (0.495, 0.695) | 0.596 (0.560, 0.631) |
| Neural late fusion | 0.685 (0.568, 0.784) | 0.614 (0.534, 0.687) | 0.594 (0.488, 0.691) | 0.584 (0.533, 0.634) | 0.586 (0.530, 0.640) | 0.585 (0.490, 0.679) | 0.597 (0.567, 0.627) |
| PriorityLasso | 0.657 (0.545, 0.759) | 0.586 (0.505, 0.659) | 0.600 (0.508, 0.688) | 0.575 (0.524, 0.628) | 0.551 (0.500, 0.599) | 0.627 (0.528, 0.724) | 0.582 (0.547, 0.617) |
| *Adapted features with TCGA scaling* | | | | | | | |
| PoE-VAE | 0.691 (0.568, 0.791) | 0.607 (0.514, 0.688) | 0.595 (0.488, 0.696) | 0.512 (0.462, 0.562) | 0.567 (0.510, 0.626) | 0.603 (0.503, 0.697) | 0.583 (0.523, 0.644) |
| Borrowing model | 0.693 (0.568, 0.794) | 0.590 (0.484, 0.686) | 0.628 (0.540, 0.720) | 0.524 (0.478, 0.570) | 0.579 (0.538, 0.617) | 0.602 (0.509, 0.696) | 0.588 (0.531, 0.645) |
| Elastic-net late fusion | 0.687 (0.566, 0.794) | 0.638 (0.555, 0.712) | 0.582 (0.480, 0.683) | 0.560 (0.511, 0.606) | 0.577 (0.520, 0.631) | 0.605 (0.507, 0.704) | 0.593 (0.549, 0.636) |
| Neural late fusion | 0.683 (0.580, 0.775) | 0.633 (0.553, 0.706) | 0.562 (0.452, 0.675) | 0.568 (0.519, 0.616) | 0.586 (0.528, 0.645) | 0.601 (0.502, 0.693) | 0.597 (0.555, 0.639) |
| PriorityLasso | 0.658 (0.554, 0.757) | 0.579 (0.500, 0.654) | 0.594 (0.501, 0.683) | 0.562 (0.510, 0.615) | 0.555 (0.504, 0.602) | 0.630 (0.530, 0.729) | 0.578 (0.542, 0.614) |
| *Fully frozen pipeline* | | | | | | | |
| PoE-VAE | 0.690 (0.567, 0.790) | 0.613 (0.523, 0.692) | 0.586 (0.478, 0.691) | 0.510 (0.458, 0.562) | 0.571 (0.514, 0.629) | 0.601 (0.499, 0.696) | 0.584 (0.523, 0.645) |
| Borrowing model | 0.690 (0.567, 0.793) | 0.598 (0.498, 0.693) | 0.579 (0.495, 0.663) | 0.519 (0.469, 0.569) | 0.610 (0.564, 0.655) | 0.604 (0.502, 0.706) | 0.591 (0.534, 0.647) |
| Elastic-net late fusion | 0.692 (0.569, 0.797) | 0.649 (0.571, 0.719) | 0.577 (0.476, 0.675) | 0.564 (0.515, 0.612) | 0.579 (0.522, 0.633) | 0.603 (0.505, 0.700) | 0.598 (0.552, 0.645) |
| Neural late fusion | 0.681 (0.577, 0.773) | 0.636 (0.557, 0.708) | 0.564 (0.453, 0.675) | 0.569 (0.521, 0.618) | 0.591 (0.533, 0.649) | 0.599 (0.500, 0.691) | 0.599 (0.558, 0.639) |
| PriorityLasso | 0.657 (0.553, 0.756) | 0.605 (0.528, 0.675) | 0.590 (0.500, 0.680) | 0.562 (0.510, 0.615) | 0.550 (0.488, 0.605) | 0.630 (0.530, 0.729) | 0.583 (0.544, 0.622) |

| (c) Contrast | Level | Cohort-adapted pipeline | Fully frozen pipeline | Adapted features with TCGA scaling (95%) |
|---|---|---|---|---|
| PoE-VAE − elastic-net late fusion | 98.33% | −0.0090 (−0.0329, +0.0148) | −0.0202 (−0.0541, +0.0137) | −0.0176 (−0.0409, +0.0058) |
| PoE-VAE − neural late fusion | 98.33% | −0.0093 (−0.0355, +0.0169) | −0.0235 (−0.0633, +0.0163) | −0.0214 (−0.0518, +0.0090) |

| (c) Contrast | Level | Cohort-adapted pipeline | Fully frozen pipeline | Adapted features with TCGA scaling (95%) |
|---|---|---|---|---|
| PoE-VAE − PriorityLasso | 98.33% | +0.0055 (−0.0490, +0.0599) | −0.0044 (−0.0553, +0.0466) | −0.0013 (−0.0376, +0.0351) |
| Borrowing model − PoE-VAE | 95% | +0.0070 (−0.0008, +0.0147) | +0.0076 (+0.0001, +0.0152) | – |
| Elastic-net late fusion − PriorityLasso | 95% | +0.0165 (−0.0124, +0.0453) | +0.0119 (−0.0125, +0.0363) | – |
| Neural late fusion − elastic-net late fusion | 95% | +0.0021 (−0.0103, +0.0145) | +0.0015 (−0.0096, +0.0127) | – |

#### Supplementary Figure S1 Late fusion and missing blocks (post hoc)

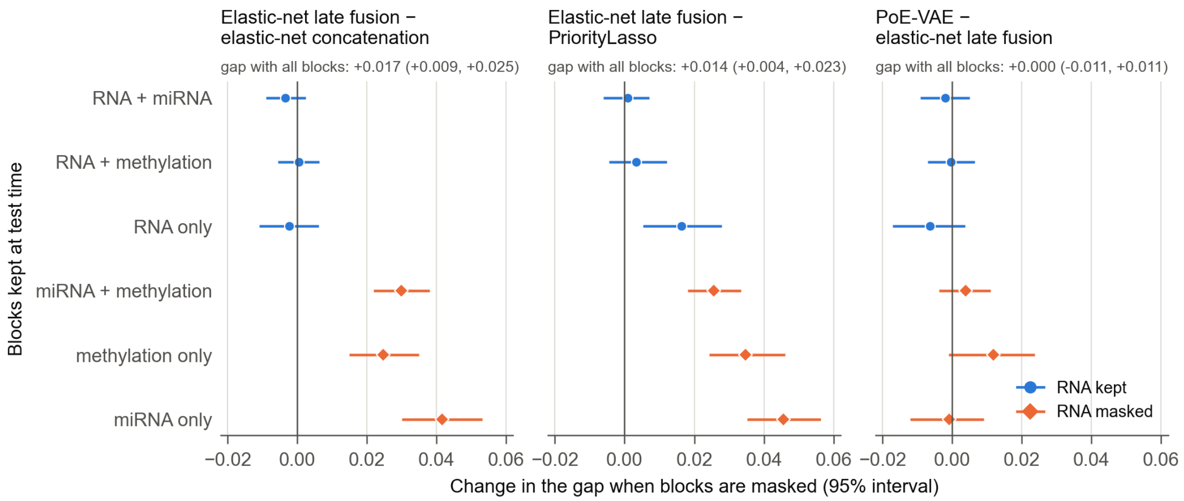


Change in the concordance gap between two models (first named minus second) after test-time masking, relative to their gap with all three blocks observed (given above each panel). Positive values indicate a larger advantage for the first-named model after masking; the plotted values are changes in the gap, not the absolute gap under masking. Test patients with all three blocks measured in the 11 three-block cancers, disease-specific survival; equal-weight means over cancers with 95% percentile bootstrap intervals. Circles: RNA kept. Diamonds: RNA masked. Elastic-net late fusion led concatenation by 0.016 (95% interval +0.008 to +0.024) under natural availability and by 0.017 (+0.009 to +0.025) with all blocks observed; masking RNA widened the gap by 0.025 to 0.042, masking microRNA or methylation produced no detected change, and PriorityLasso showed a similar pattern. miRNA, microRNA.

#### Supplementary Figure S2 External evaluation

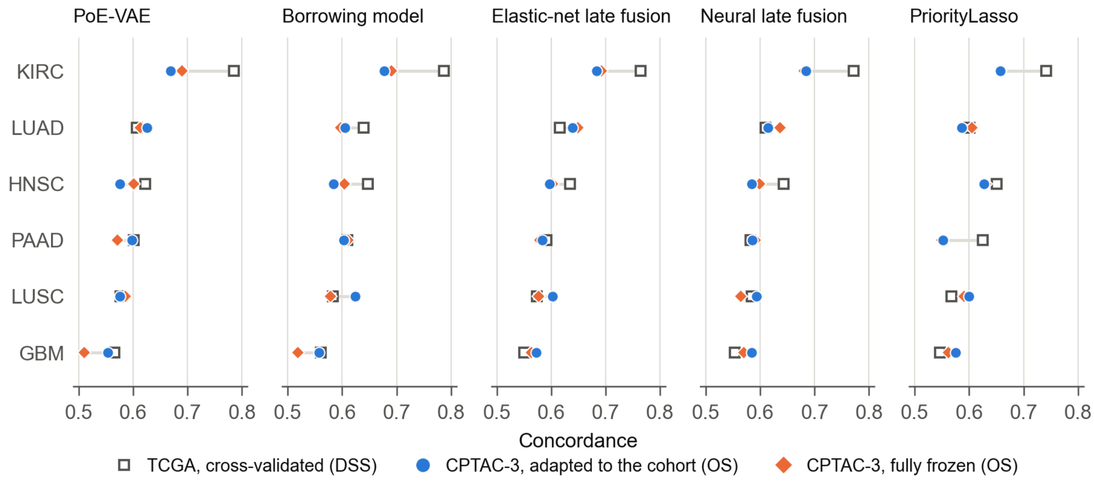


Concordance in six CPTAC-3 cohorts (overall survival) under the cohort-adapted pipeline (circles) and the fully frozen pipeline (diamonds), beside the cross-validated concordance of the same model in TCGA (disease-specific survival; open squares). The cohort-adapted pipeline selects features by their coverage and variability in the external cohort and scales them with the external cohort's own statistics; the fully frozen pipeline retains the full TCGA feature set, uses TCGA scaling statistics and imputes missing external values with TCGA training medians. Internal and external values differ in endpoint, feature availability and training size and are compared descriptively. Per-cohort intervals and paired contrasts are in Supplementary Table S7.

### Supplementary notes

#### Supplementary Note 1. Studies of cross-cancer survival learning and their released code

The table lists twelve selected peer-reviewed studies that report survival prediction with a model that uses data from other cancers, in most cases beside a model that does not. The selection was not systematic and is not a representative sample of the literature, so it supports no statement about how common a practice is. The comparisons estimate different quantities and are not combined. Entries record what the cited main reports state; a dash means that we did not record a single paired difference in concordance for the study. Endpoint and target training size are given for the two studies whose gains are quoted in the main text [10, 11].

| Study | Type of comparison | Reference models and design | Reported result | Scope of the comparison |
|---|---|---|---|---|
| VAECox; Kim et al. 2020 [10] | Transfer: gene-expression VAE pretrained on 20 cancers, then cancer-specific survival fitting | Cox-LASSO, Cox-ridge and Cox-nnet references; ten 80/20 splits with five-fold tuning on training data. Endpoint: time to death. Target training set: 80% of each cancer's patients | Micro-average gain of 0.020 from transfer plus fine-tuning (Methods 2.5, Results 3.1-3.2) | Repeated splits and penalised Cox comparators are included. The 0.020 is distinct from gains over other model families. |
| Song et al. 2026 [11] | Joint training: shared encoders with cancer-specific heads; histology, RNA and clinical text | Cancer-specific training; five random seeds and paired tests. Endpoint: time from diagnosis to death. Target training set: 40% of each cancer's patients (4:2:4 training, validation and test split); ten cancers with more than 150 patients each | Mean concordance 0.670 against 0.625 (Results on cross-cancer training and statistical validation; Conclusion) | Joint learning with target training patients included, not target-excluded pretraining. |
| TMO-Net; Wang et al. 2024 [9] | Pretraining: pan-cancer multi-omics representation, then fine-tuning | VAECox, SNN and DeepSurv references; the Methods describe five-fold procedures and fixed random seeds | – | A fixed seed with folds does not establish five independently repeated partitions. The main text alone does not establish patient-level overlap between each survival test set and pretraining. The released code (below) keeps the highest test-fold concordance over fine-tuning epochs and includes the target cancer's training patients in each fold's pretraining. |
| TMO-Net+; Liu et al. 2026 [13] | Pretraining: supervised pan-cancer pretraining, then task-specific fine-tuning | PCLSurv, MMOSurv and TMO-Net; five seeds; its Table 4 gives concordance intervals for individual models | – | A random-initialisation control is described for metastasis prediction, not for the survival table. Individual-model intervals are not paired intervals for the benefit of pretraining. |
| SeNMo; Waqas et al. 2025 [14] | Joint training: pan-cancer fitting | 80% training-validation and 20% held-out data within each cancer; ten-fold checkpoints averaged and ensembled on the held-out set | – | Ten checkpoints do not imply ten independently resampled held-out sets. The comparison concerns joint fitting and tumour-specific application or fine-tuning. |

| Study | Type of comparison | Reference models and design | Reported result | Scope of the comparison |
|---|---|---|---|---|
| Qiu et al. 2020 [7] | Few-shot adaptation: meta-learning with the target cancer excluded from auxiliary training | Direct learning with 20, 150 and 250 target patients on common test cases; adaptation uses 20 patients; five-fold tuning on glioma | – | A gain over 20-patient direct learning does not establish the same gain over a full-size target model. |
| MMOSurv; Wen and Li 2025 [15] | Few-shot adaptation: multi-omics meta-learning with an independent target cancer | Direct learning, multitask learning and regular pretraining; 10 or 20 target training samples; 50 target draws and 20 repetitions | – | Hyperparameters are searched in the 20-shot setting and reused at 10 shots. The main text does not identify which data were used to select them. |
| Cheerla and Gevaert 2019 [16] | Joint training: multimodal pan-cancer learning | Cancer-specific models; an 85/15 split with the same test cases for the per-cancer comparison | KIRC concordance 0.73 with pan-cancer training and 0.78 with cancer-specific training (its Table 3) | Target training patients are included in joint learning. |
| Transfer-Cox; Li et al. 2016 [17] | Transfer: source and target Cox models share sparsity through an L2,1 penalty | Local and global Cox, LASSO-Cox and elastic-net Cox; five-fold evaluation | – | Target-only penalised Cox references are present. Its Table III reports per-model standard deviations, not an interval on the paired transfer difference. |
| Wang et al. 2017 [18] | Joint training: multitask Cox models on 21 TCGA cancers using microRNA data | Individual-cancer and global Cox, LASSO-Cox and elastic-net Cox | – | Concordance and standard deviations are reported per model. Joint learning differs from a pretrained representation fitted without the target cancer. |
| Samorodnitsky et al. 2022 [19] | Joint model: hierarchical spike-and-slab survival regression on BIDIFAC+ components across 29 cancers | Shared, joint and separate-cancer models; five-fold mean log-posterior predictive likelihood | Not a concordance measure | Its gain cannot be interpreted as a concordance gain. |
| Sabbatini et al. 2026 [8] | Transfer: leave-one-cancer-out pretraining, then fine-tuning; RNA, 27 cancers | Tumour-specific neural models; random 70/30 splits of the target cohort (50 per reference configuration and 200 per transfer configuration in the released notebooks) | – | The released notebooks record the held-out concordance of every candidate configuration and have no validation set separate from the held-out part; the step that chooses a configuration is not in the released code (below). Excluding the target cancer from pretraining does not by itself make the choice of configuration independent of the held-out data. |

**Released code.** The following paragraphs record what the released code does at the stated commits. We did not run either code base.

**TMO-Net.** We read the TMO-Net repository (github.com/FengAoWang/TMO-Net) at the last commit before publication (50c890fd, 23 May 2024; the article was published on 6 June 2024) and at the current commit (1ee4ef8b, 23 December 2025), together with the TCGA archive in Zenodo record 11258239. Statements hold at both commits unless stated otherwise.

Test-fold selection. In train/train_tcga_pancancer_multitask.py, cal_c_index() scores the test fold after every fine-tuning epoch and keeps the highest concordance (lines 466 to 468 at 50c890fd; lines 332 to 334 at 1ee4ef8b), which is written as the result for that cancer and fold. There is no validation split. The pretrained and randomly initialised arms share this procedure.

Target cancer in pretraining. TCGA_Dataset_pretrain() builds each fold's pretraining set from the fold-assignment file without a cancer filter, and CancerDataset then uses every cancer type. Survival fine-tuning for a target cancer uses the same fold and that fold's pretrained model. The target's test-fold patients are therefore excluded from pretraining and its training-fold patients are included.

Released inputs. The code reads harmonised omics files, a cleaned clinical table and fold-assignment files (train_data.csv and test_data.csv). The Zenodo archive provides the omics matrices and a clinical table but not these files, so the published folds cannot be reconstructed.

**Sabbatini et al. [8].** We read the notebooks released with the study (gitlabdigei.aizoon.it/aizoon/survival-transfer-learning, commit 990709e2, 9 December 2025) without running them. The baseline notebook samples 750 configurations at random and, for each, fits the network on 50 random 70/30 splits of the target cohort, recording the concordance in the training part and in the held-out part; features are min-max scaled over all patients of the cohort before the splits are drawn. The transfer notebook loads a saved pretrained network and, for each of 96 fine-tuning configurations, refits it on 200 random 70/30 splits of the target cohort and records the same two values. Neither notebook has a validation set separate from the held-out part, and the step that chooses a configuration is not in the released code. The notebooks require the DeepHazard package, whose licence must be requested from its owner, so we did not rerun the study.

#### Supplementary Note 2. Models and evaluation methods

**PoE-VAE.** The model is a product-of-experts variational autoencoder [23, 24] with a proportional-hazards survival head [25]. Each omics block has an encoder with one hidden layer of 128 units and layer normalisation [26], which produces a diagonal Gaussian posterior over a 48-dimensional latent variable. The posteriors for the observed blocks are combined with a standard normal prior using a product of experts [24], yielding a single Gaussian distribution over the shared latent representation. Let $\boldsymbol{\mu}$ denote its mean vector and $\sigma_k$ its standard deviation in latent coordinate $k$. A linear projection with coefficient vector $\mathbf{w}$ gives a Gaussian log-risk $\eta$, with mean $\mathbf{w}^{\mathrm{T}}\boldsymbol{\mu}$ and variance $\sum_{k=1}^{48} w_k^2 \sigma_k^2$. The survival model is proportional hazards,

$$h(t \mid \eta) = h_0(t) \exp(\eta)$$

where $h_0(t)$ is a piecewise-exponential baseline hazard [27], with knots at the quartiles of the training event times and learned rates. Its cumulative baseline hazard is $H_0(t) = \int_0^t h_0(u)\mathrm{d}u$. For a patient with observed follow-up time $t$ and event indicator $\delta$, the survival loss is

$$- \log \mathrm{E}_{\eta} \left[ (h_0(t) e^{\eta})^{\delta} \exp\left(-H_0(t) e^{\eta}\right) \right]$$

where the expectation over the Gaussian log-risk is computed by 64-node Gauss–Hermite quadrature [28]. The total loss also includes 0.1 times the mean squared reconstruction error across observed blocks, computed from a latent sample using one decoder per block, and 0.001 times the Kullback–Leibler divergence [29] from the standard normal prior, averaged over latent coordinates. At each step, the encoders receive a subset sampled uniformly from the non-empty subsets of each patient's observed blocks, while reconstruction targets all observed blocks, following subset training in multimodal variational autoencoders [24]. Training used 100 full-batch steps of AdamW [30], with a learning rate of 0.001, weight decay of 0.0001 and gradient norm clipped at 5. Concordance was computed using the mean log-risk.

Each encoder is Linear(features, 128), layer normalisation, GELU and Linear(128, 96), which gives the 48 means and 48 raw log-variances of the block's posterior; each log-variance is bounded as 6 tanh(raw / 6). Each decoder is Linear(48, 128), layer normalisation, GELU and Linear(128, features). The reconstruction error is the squared error averaged over the features of a block, then over the observed blocks of a patient, then over patients. Numerical integration and initialisation follow the released code.

**Shared feature set.** Within each non-target cohort, features were required to have no more than 20% missing values and at least two finite measurements among primary-tumour patients with the corresponding assay. RNA features were restricted to the supplied Ensembl protein-coding gene list and required positive expression in at least 10% of these patients; microRNA features required positive expression in at least 20%. Methylation probes were restricted to chromosomes 1-22 and required an empty Probe_SNPs_10 annotation in the Illumina HumanMethylation450 manifest. Candidate features had to satisfy these criteria in every non-target cohort with an eligible block. Within each non-target cohort, missing values were median-imputed before feature-wise median absolute deviations were calculated, using M-values for methylation; features were then ranked by the mean of these deviations across cohorts (Methods 2.2).

**Variants of the borrowing analysis.** The reference analysis selects features in the 19 non-target cohorts, excludes all target-cancer patients from pretraining and selects the fusion weight by inner cross-validation. Five further analyses change the features, pretraining data, selection procedure or training scheme (Supplementary Table S4). These variants do not form a one-factor-at-a-time decomposition.

Target-selected features. Before partitioning, the shared feature-eligibility criteria described above were applied to all target-cancer patients with the relevant block. Methylation probes were also screened to retain the 20,000 with the largest median absolute deviations across those patients. Both steps used omics values without outcomes, but included patients subsequently used for testing. Within each training partition, missing values were filled with training medians, and features were ranked by median absolute deviation among training patients with the block. The top 4,000 genes, all eligible microRNAs and the top 4,000 methylation probes were retained and standardised on the training patients. This comparison therefore evaluates a pipeline with screening before partitioning; feature selection was not confined entirely to training patients.

Target-selected features gave no detected gain for the PoE-VAE (−0.0001, 97.5% interval −0.014 to +0.014) or elastic-net late fusion (+0.002, −0.007 to +0.012; Supplementary Table S4).

Target-inclusive pretraining. The representation was pretrained on the 19 other cohorts and all target-cancer patients, including test patients, without their outcomes.

Test-selected weight. For each cancer, the fusion weight was selected from (0, 0.1, . . . , 1) to maximise mean test concordance across its 25 partitions.

Best of three borrowing models. For each cancer, we selected the model with the highest mean test concordance among the Cox probe on the pretrained representation alone, fusion with a fixed weight of 0.5 and the borrowing model.

Joint training. One model was trained with outcomes from the 19 other cohorts and the target cancer's training patients, using a separate baseline hazard for each cohort. The shared risk projection then scored target-cancer test patients without further target-specific fitting.

**Elastic net.** Elastic-net fitting [33] used a path of 30 penalties with mixing parameter 0.5, an initial minimum-to-maximum penalty ratio of 0.05 and retries at 0.1 and then 0.2 after a numerical failure of the full-training path. Inner folds with a numerical failure or no validation event were omitted, with at least three usable inner folds required, and the chosen penalty was evaluated from the fitted full-training path. In the 3,270 retained final fits, no fallback path, failed inner fold or event-free validation fold occurred, and every fit used all five inner folds.

**PriorityLasso.** The R package prioritylasso version 0.4.0 contains an indexing expression in calculate_offsets() that can reduce a matrix to a vector when exactly one row is selected. The expression complete.cases(X[current_missings, blocks[[i]]]) was replaced with complete.cases(X[current_missings, blocks[[i]], drop = FALSE]). This preserves matrix dimensions for the completeness check. The wrapper requires version 0.4.0 and exactly one matching expression before applying the change; it stops if either condition fails. The correction changes neither the model's penalty nor the block-order rule. The first block is penalised, each penalty uses minimum five-fold cross-validated deviance, and block order uses training-only cross-validated single-block concordance. Missing training blocks use offset imputation; missing test blocks use the package's zero-block rule.

**Clinical model.** The clinical model is a ridge Cox model [31] with the penalty fixed at 0.1. The eligibility of each variable was determined from counts over all eligible patients of the cancer, without outcomes, by the rules in the legend of Supplementary Table S1. Stage and grade were coded as integers. Missing values were imputed with training medians, missingness indicators for stage and grade were added when missing values occurred in training, constant training columns were removed and the remaining columns were standardised using training means and standard deviations.

**VAECox reimplementation.** We reimplemented VAECox from its public repository (github.com/dmis-lab/VAECox, commit b9946773, 13 July 2020). Architecture: encoder Linear(genes, 4,096) with tanh; mean and log-variance heads Linear(4,096, 128) with tanh; decoder Linear(128, 4,096), tanh, Linear(4,096, genes), applied to the mean as in the published code. Pretraining loss: mean squared reconstruction error plus the Kullback–Leibler divergence averaged over patients and latent coordinates plus $10^{-6}$ times the L1 norm of all parameters; Adam with learning rate 0.001 and weight decay $10^{-5}$, 501 full-batch steps, final model.

Survival network. The pretrained encoder and mean head were followed by a Cox-nnet head: Linear(128, 12), tanh, Linear(12, 1). All weights were fine-tuned in double precision under the Breslow partial likelihood, using Adam for 21 full-batch steps.

Hyperparameter search. The grid combined learning rates of $10^{-4}$ or $10^{-3}$ with weight decay of $10^{-3}$, $10^{-4}$ or $10^{-5}$. The published grid also varies a dropout rate that the VAECox network does not use. Unstratified inner five-fold cross-validation scored each candidate by its highest mean validation concordance across steps. A later candidate replaced an earlier one only if its score was strictly higher.

Evaluation step. Fine-tuning was fixed at 21 steps, and the target-excluded protocol evaluated each network at the last step. Thus, candidates were selected by their best validation step but evaluated at a potentially different step. A follow-up also evaluated the target-excluded protocol at the chosen candidate's best mean inner validation step, selected separately for each network and partition (Supplementary Table S5d).

Differences from the original study. RNA from each target's 4,000-gene feature set rather than 20,502 imputed genes; disease-specific survival rather than time to death; our 20 cohorts as the pretraining pool; five repeats of five-fold cross-validation rather than ten 80/20 splits; per-cohort z-scoring (the repository reads stored values without scaling); the 80% pretraining sample drawn with a recorded seed; and inner folds without a comparable pair dropped. The repository has no code for the randomly initialised reference reported in the paper; ours keeps the VAE's initial weights. The reimplementation is therefore a test of the published method and evaluation procedure on these data, not a rerun of the original experiment.

Within each cancer and seed, every VAE (published, test-excluded and target-excluded pools) started from the same initial weights, and the randomly initialised encoder was that untrained VAE. The Cox-nnet head had the same initial weights in every version, and all versions shared inner folds and test patients. Pretraining reduced the loss from 1.08 to between 0.51 and 0.68 in every cancer. The pretraining pools nested as intended: 6,254 patients for the published pool, fewer for the test-excluded pool and fewest for the target-excluded pool. The fine-tuned networks reached concordance similar to the other models (0.65 on average for the randomly initialised network).

Ten-cancer overall-survival setting. Targets: BLCA, BRCA, HNSC, KIRC, LGG, LIHC, LUAD, LUSC, OV and STAD. Pretraining cancers: these ten plus UCEC, THCA, SKCM, PRAD, PAAD, LAML, KIRP, GBM, COAD and CESC (8,865 patients with RNA data). RNA for THCA, SKCM, PRAD and LAML came from the same GDC Xena hub as the other cohorts. Samples were primary tumours except primary blood for LAML and, for SKCM, a primary tumour where available and otherwise a metastasis, because most SKCM samples are metastases. Genes: protein-coding genes present in all 20 cohorts with at most 20% missing values and positive expression in at least 10% of the pooled samples (18,858). Endpoint: overall survival (365 to 1,080 eligible patients and 125 to 261 deaths per target). Values were checked against an independent reader in every cohort, and the 16 cohorts shared with the main analysis selected the same patients from the same files. Pretraining reduced the loss from 1.09 to between 0.71 and 0.79 in every cancer. This setting uses the target cancers, pretraining cancers and endpoint of the original study, and replaces the 4,000-gene sets with a pooled set of 18,858 genes, against 20,502 imputed genes in the original. It still differs from the original in expression processing (the original used imputed data that we did not have), in resampling and in the reconstructed randomly initialised reference.

**Survival probabilities.** We converted risk scores into survival probabilities by fitting a free Cox slope and Breslow baseline [25, 35, 38] to out-of-fold training risks. The fitted mapping was applied to outer test risks standardised using the outer model's training-risk moments. BlockForest supplied its own curves; the PoE-VAE's learned baseline was not used. The Brier comparison therefore evaluates each complete probability pipeline, including its mapping.

This procedure replaced the originally specified mapping fitted to in-sample training risks. Median training concordance was 0.99 for the PoE-VAE and 1.00 for neural late fusion, compared with 0.89 for elastic-net late fusion. Test concordance was about 0.65 for all models. Slopes fitted to the nearly separating training risks diverged and produced test survival probabilities close to 0 or 1. We changed the mapping after examining these training-data diagnostics and before computing any Brier score.

For the borrowing model, the mapping reused the fusion weight and probe penalty selected on the complete outer-training set. Inner validation outcomes had therefore contributed to those two choices, although no outer test outcome entered the mapping.

The Brier grid [40] used all eligible event times within a cancer to define its 10th and 80th percentiles, while censoring weights came from each training fold. Thus the evaluation horizon was outcome-dependent, shared across arms, and separate from model fitting. Brier scores were averaged by patient within repeat, whereas concordance was averaged by fold. These different weights can contribute to different rankings. Bootstrap intervals condition on the fitted predictions, mappings and censoring weights.

**Nesting of preprocessing and selection.** In the reference borrowing analysis, feature imputation and standardisation were fitted on the outer-training set. The probe penalty was also selected on that set by inner cross-validation. These choices were reused during inner cross-validation for fusion-weight selection. Within each inner fold, the PoE-VAE and probe were refitted on the inner-training patients, as was the standardisation of the pretrained representation.

For the tuned models, inner folds were formed after outer-training imputation and scaling. In both cases, the inner scores therefore do not evaluate a pipeline refitted entirely within each inner fold. These reference and tuning procedures used no outer test data. The borrowing variants described above deliberately departed from this design: target-selected features used outcome-free screening that included future test patients, target-inclusive pretraining included test patients without outcomes, and test-selected weights and the best of three borrowing models used test concordance for selection.

**Intervals.** Bootstrap intervals describe uncertainty from sampling test patients conditional on the fitted models. They exclude the additional variation that would arise from refitting models on new patients.

Run-variation sensitivity. Before pooling, we added the sample variance of each cancer's five run-level differences, divided by five, to its bootstrap variance. This was a heuristic allowance for run-to-run variation among the same patients, not an estimate of variation from a new training cohort. It gave a borrowing estimate of +0.0023 (−0.0020 to +0.0067).

Modified intervals. With the modified Hartung–Knapp method [46], conclusions were unchanged for borrowing, the PoE-VAE comparisons with both elastic-net models, neural late fusion and PriorityLasso, and the site-held-out analyses. Two descriptive protocol contrasts lost their positive lower bounds (Supplementary Table S4). The modification was not applied to the BlockForest, clinical, survival-probability or external analyses.

Zero bootstrap variance. A cancer with zero variance was assigned the smallest positive variance observed for that contrast. This was required only for the best of three borrowing models against the borrowing model, in BLCA, GBM, KIRC and LGG. In those cancers, the borrowing model was itself the best of the three, giving an exactly zero difference (Supplementary Table S4).

#### Supplementary Note 3. Development and analysis chronology

**Analysis history.** The study developed in stages, each with a written specification before it was run, apart from the post hoc analyses listed below; later specifications were informed by earlier results from the same patients (table below). The interval adjustments apply to the named contrasts within a family; they do not correct for every analysis in the paper or for model development.

**Development.** The borrowing configuration was chosen through exploratory work on overall survival in KIRC and HNSC, both of which are among the 13 targets. This work compared supervised and unsupervised pretraining, probes on frozen encoders, fine-tuning from a fitted probe, related-cancer pretraining pools and fusion rules.

Each development stage had a written rule for proceeding. The first required a pooled gain of at least 0.02 over the reference, with at least four of five seeds positive in both cancers. No configuration met its rule. Fusion with a cross-validated weight gave a pooled gain of +0.006 (95% interval −0.007 to +0.019) in the two cancers and was taken forward for testing in all 13 cancers. The main analysis therefore tests a configuration that had not shown a gain during development.

Retaining 25% or 50% of the training patients did not detectably change the borrowing gain. Under masked evaluation, the quarter-versus-full difference was −0.0037 (95% interval −0.0136 to +0.0062), and the half-versus-full difference was +0.003 (−0.003 to +0.010). Under natural availability, the quarter-versus-full difference was −0.0003 (−0.0111 to +0.0110). These analyses were exploratory and are not independent of the main analysis.

**Post hoc analyses.** The arithmetic mean across cancers, direct contrasts between protocol variants, single-run variation of the borrowing gain, test-time masking, the endpoint sensitivity analysis (Supplementary Table S4) and the Brier contrast between the PoE-VAE and elastic-net concatenation were added after the corresponding results had been inspected. The unequal test-step selection experiment and the single-run variation of the model comparisons (Supplementary Tables S5 and S6) were specified on 10 October 2026, after their point estimates had been computed from stored summaries; they use stored predictions and refit nothing.

Analysis families and when each was specified. Retrospective account compiled from the dated written specifications of the study. Each family down to the ten-cancer overall-survival setting, and the validation-selected step, was specified in writing before it was run; the summaries and sensitivity analyses, unequal test-step selection and the single-run variation of the model comparisons were specified after their results had been inspected. Later families were specified with knowledge of earlier results from the same patients. The interval levels were therefore adjusted for the named contrasts within a family only and carry no guarantee across the study, and "prespecified" in the manuscript refers to this sequence, not to a registered protocol. Contrasts are first-named minus second-named. Unless otherwise stated, internal analyses used disease-specific survival in the 13 TCGA cancers under natural block availability; levels apply to the random-effects summary.

| **Analysis** | **Contrasts** | **Interval** | **Specified** | **Status** |
|---|---|---|---|---|
| Borrowing | Borrowing model − PoE-VAE | 95% | Before its results, after exploratory model development and endpoint selection | Primary |
| Elastic-net references | PoE-VAE − elastic-net late fusion; PoE-VAE − elastic-net concatenation | 97.5% | After the borrowing results | Two co-primary contrasts |
| Feature selection | Target-selected − shared features, for the PoE-VAE and for elastic-net late fusion | 97.5% | After the elastic-net results | Two co-primary contrasts; no detected gain, and the shared feature set was retained |
| PriorityLasso and neural late fusion | PoE-VAE − PriorityLasso; PoE-VAE − neural late fusion | 97.5% | After the feature-selection result, which fixed the feature set | Two co-primary contrasts |
| Site-held-out folds | Site-held-out − random folds, for the PoE-VAE and for elastic-net late fusion | 97.5% | After the random-fold results | Two co-primary contrasts |
| Departures from the protocol | Joint training, target-inclusive pretraining, test-selected weight and best of three borrowing models, each − PoE-VAE; target-inclusive pretraining − borrowing model | 95% | Before their results | Descriptive, without multiplicity correction. Intervals for test-selected quantities condition on the selection |
| BlockForest | PoE-VAE − BlockForest; elastic-net late fusion − BlockForest | 97.5% | Before its results | Two co-primary contrasts |
| Clinical model | PoE-VAE − clinical model; elastic-net late fusion − clinical model | 97.5% | Before its results | Interpreted by the sign of the interval |
| External cohorts, cohort-adapted pipeline | PoE-VAE − elastic-net late fusion, neural late fusion and PriorityLasso | 98.33% | Before its results; scaling with TCGA statistics was a prespecified sensitivity analysis | Primary external analysis; six CPTAC-3 cohorts, overall survival |
| External cohorts, fully frozen pipeline | As above | 98.33% | After the cohort-adapted results, before it was run | Exploratory |
| VAECox reimplementation | Pretrained − randomly initialised, target-excluded protocol | 95% | 8 October 2026, after every analysis above was reported and before its models were fitted | Primary for this analysis; the other versions and their differences are descriptive |

| Analysis | Contrasts | Interval | Specified | Status |
|---|---|---|---|---|
| Tuned models | Tuned PoE-VAE − elastic-net late fusion; tuned neural late fusion − elastic-net late fusion | 97.5% | As for the VAECox reimplementation | Two co-primary contrasts |
| VAECox, ten-cancer overall-survival setting | Pretrained − randomly initialised, target-excluded protocol | 95% | 9 October 2026, after the first VAECox results | Exploratory; ten cancers, overall survival |
| Summaries and sensitivity analyses | Arithmetic mean across cancers; modified Hartung–Knapp intervals; direct contrasts between protocol variants; single-run variation of the borrowing gain; test-time masking; endpoint sensitivity; Brier contrast with elastic-net concatenation | As stated | After the corresponding results were inspected | Post hoc |
| Unequal test-step selection; single-run variation of the model comparisons | Supplementary Tables S5 and S6 | 95% | 10 October 2026, after their point estimates had been inspected | Post hoc |
| VAECox, validation-selected step | Pretrained − randomly initialised, target-excluded protocol, in both settings | 95% | 12 October 2026, after every earlier VAECox result was known and before it was run | Exploratory |

Table 3 of the main manuscript combines comparisons from the elastic-net, PriorityLasso and neural late fusion, BlockForest and tuned-model families. Its 97.5% intervals were assigned within these families, not jointly over all rows. Uno's concordance is secondary at the corresponding contrast's level, and integrated Brier score comparisons use nominal 95% intervals.
